# High-Throughput and High-Dimensional Single Cell Analysis of Antigen-Specific CD8^+^ T cells

**DOI:** 10.1101/2021.03.04.433914

**Authors:** Ke-Yue Ma, Alexandra A. Schonnesen, Chenfeng He, Amanda Y. Xia, Eric Sun, Eunise Chen, Katherine R Sebastian, Robert Balderas, Mrinalini Kulkarni-Date, Ning Jiang

**Affiliations:** Interdisciplinary Life Sciences Graduate Programs, the University of Texas at Austin, Austin, Texas, USA; Beijing Advanced Innovation Center for Biomedical Engineering, Beijing, China; Department of Biomedical Engineering, the University of Texas at Austin, Austin, Texas, USA; Department of Molecular Biosciences, the University of Texas at Austin, Austin, Texas, USA; Department of Nutritional Sciences, the University of Texas at Austin. Austin, Texas, USA; Department of Internal Medicine, Dell Medical School, the University of Texas at Austin, Austin, Texas, USA; Becton Dickinson, San Jose, California, USA; Department of Oncology, Dell Medical School, the University of Texas at Austin, Austin, Texas, USA

## Abstract

Although critical to T cell function, antigen specificity is often omitted in high-throughput multi-omics based T cell profiling due to technical challenges. We describe a high-dimensional, tetramer-associated T cell receptor sequencing (TetTCR-SeqHD) method to simultaneously profile TCR sequences, cognate antigen specificities, targeted gene-expression, and surface-protein expression from tens of thousands of single cells. Using polyclonal CD8^+^ T cells with known antigen specificity and TCR sequences, we demonstrated over 98% precision for detecting the correct antigen specificity. We also evaluated gene-expression and phenotypic differences among antigen-specific CD8^+^ T cells and characterized phenotype signatures of influenza- and EBV-specific CD8^+^ T cells that are unique to their pathogen targets. Moreover, with the high-throughput capacity of profiling hundreds of antigens simultaneously, we applied TetTCR-SeqHD to identify antigens that preferentially enrich cognate CD8^+^ T cells in type 1 diabetes patients compared to healthy controls, and discovered a TCR that cross reacts between diabetic and microbiome antigens. TetTCR-SeqHD is a powerful approach for profiling T cell responses.

## Introduction

Due to their multi-facet role in controlling infection, fighting against cancer, and responding to vaccines, the T cell has been subjected to extensive analysis^1, 2^. Recently developed multi-omic single cell profiling methods have enabled multi-dimensional analysis in single T cells, such as combining ATAC-seq with single cell RNA-seq (scRNA-seq)^3^, DNA-labeled antibody based phenotyping with scRNA-seq (CITE-seq^4^ and REAP-seq^5^), and DNA-labeled antibody based phenotyping with targeted single cell gene expression^6^. They have greatly advanced our understanding of T cell immune responses in multiple disease settings^7-9^.

T cell antigen-specificity, although critical to T cell function and T cell based immunotherapy development, has been challenging to analyze in a high-throughput manner until recently. Using T cell trogocytosis^10^ or reporter genes^11-14^, a suite of technologies have been developed in this areas enabling the high-throughput screening for T cell antigens, such as SABR^11^, MCR-TCRs^12^, T-scan^13^, granzyme B based target cell tag^14^. These methods have provided much needed T cell epitope information in the context of cancer^11-14^ and SARS-CoV-2^15^. However, because these methods use expanded T cells or TCR-transduced cell lines, they do not support the profiling of phenotype or gene expression in primary T cells. Peptide-MHC tetramer (pMHC) based methods can be applied to primary T cells. In combination with mass cytometry, it has been shown that over 100 antigens can be screened in parallel along with phenotype^16, 17^. Yet, the destructive nature of mass cytometry prevents the acquisition of TCR sequences, which is critical for T cell antigen validation.

We previously developed TetTCR-Seq^18^ to link the T cell receptor (TCR) sequence information to its cognate antigens by sequencing DNA-barcoded pMHC tetramers bound on individual single T cells^18^. TetTCR-Seq enables the screening of hundreds of antigens on primary T cells. To better understand functional profiles of antigen-specific CD8^+^ T cells, a method to simultaneously profile two other “dimensions” of parameters, gene expression and surface-protein expression, is imperative. In this study, we describe a high-dimensional TetTCR-Seq (TetTCR-SeqHD) method that enables us to simultaneously profile paired TCR sequences, cognate antigen specificities, targeted gene expression, and selected surface-protein expression in tens of thousands of single cells from multiple biological samples. Using a mixture of T cell clones, we demonstrated high precision and recall rates with TetTCR-SeqHD. We then developed a panel of 215 endogenous antigens, majority of which are type 1 diabetes (T1D) related peptides, and 65 foreign antigens. Using these antigens on a set of primary CD8^+^ T cells from a cohort of healthy individuals and T1D patients, we showed that foreign pathogen-specific T cells exhibited infection dependent states. Analyzing the 209 T1D related peptides, we identified three peptides that have elevated antigen-specific CD8^+^ T cell frequencies in T1D patients compared with healthy controls. Transducing TCR sequences identified into human CD8^+^ T cells allowed us to functionally validate these TCRs including one that cross-reacts between a T1D related peptides and a peptide derived from microbiome. TetTCR-SeqHD together with the flexibility and the speed of generating high-throughput antigen libraries through *in vitro* transcription and translation (IVTT), created a powerful technology to characterize the function and phenotype and track clonal lineage of antigen-specific T cells at single cell level in one assay.

## Results

### TetTCR-SeqHD detects correct antigen specificities in polyclonal CD8^+^ T cells

In Tetramer associated TCR Sequencing High Dimension (TetTCR-SeqHD) technology, each peptide encoding DNA oligo was individually *in vitro* transcribed/translated (IVTT) to generate corresponding peptide, which was later loaded onto MHC molecules. Then peptide-MHC (pMHC) tetramer was tagged with its corresponding peptide oligo bearing a 3’ polyA overhang, which serves as the DNA barcode for that antigen specificity (**Fig. 1a**). This enables the tetramer barcodes to be captured by BD Rhapsody^™^ beads and reverse transcribed together with other mRNAs captured, including TCR transcripts (**Fig. 1b**). At the same time, 59 DNA-labeled antibodies^19^ were used to stain cells. Similar to the tetramer, the DNA barcodes labeled to the antibodies were captured by the same beads. Thus, TetTCR-SeqHD integrates TCR sequencing with TCR antigen-specificity, gene expression, and phenotyping in tens of thousands of single cells for hundreds of antigens simultaneously.

**Figure 1.**
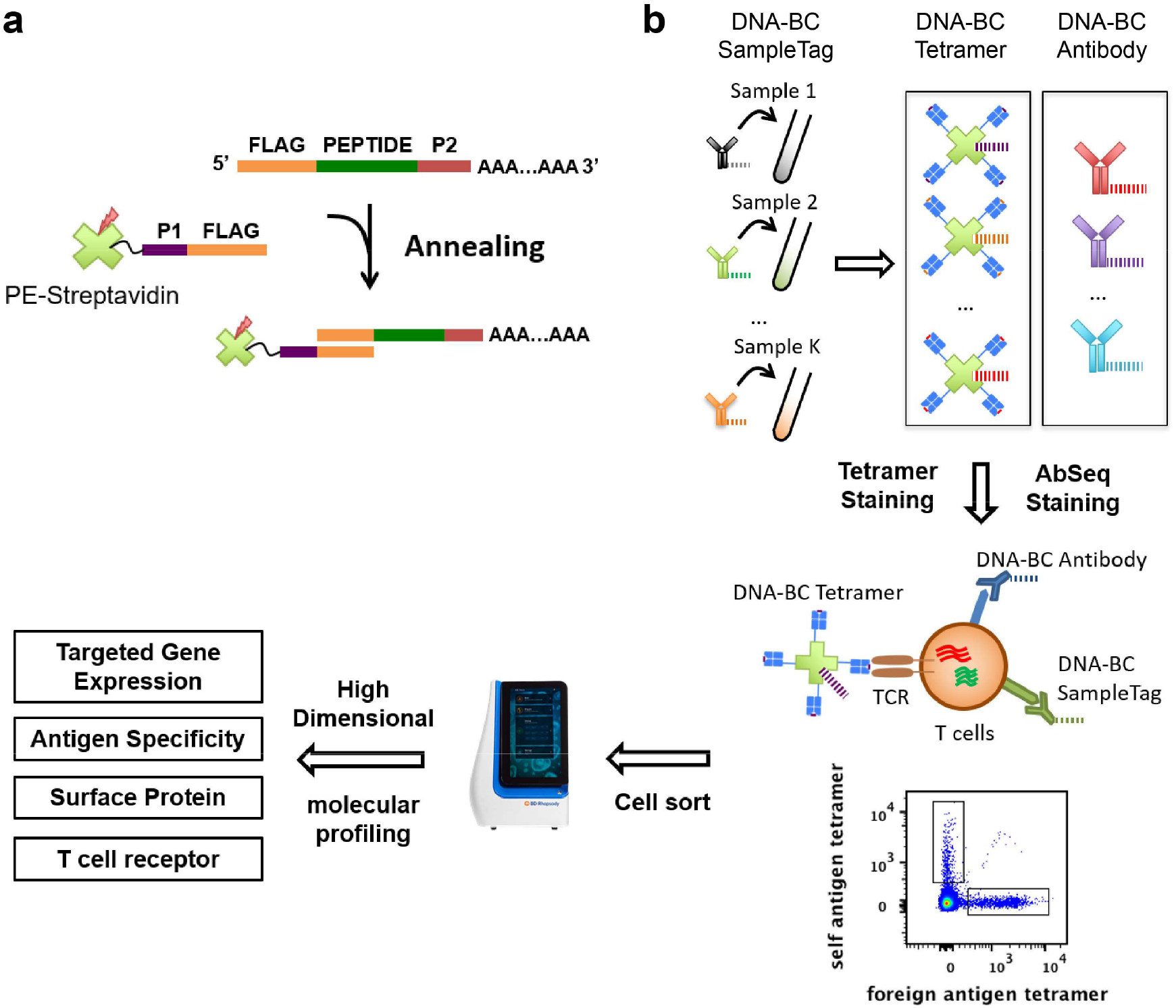
Schematics of TetTCR-SeqHD workflow. (**a**) DNA tetramer barcode encoding antigen DNA sequence was synthesized with a 3’ polyA tail. Fluorophore labeled streptavidin conjugated with an oligonucleotide sequence complementary to the 5’ end of DNA tetramer barcode was then used to anneal to each unique DNA tetramer barcode to generate barcoded streptavidin. Barcoded streptavidin was further used to form tetramers with cognate-antigen loaded MHC. (**b**) Each sample was stained with a unique DNA-barcoded anti-CD50 antibody as a SampleTag, a panel of DNA-barcoded tetramers, and a panel of 59 DNA-barcoded antibodies. Stained cells were sorted and then loaded on BD Rhapsody single sell analysis platform for high-throughput and high-dimensional molecular profiling, including TCR sequences, cognate antigen specificity, targeted gene expression, and surface protein level.

We first assessed the precision of TetTCR-SeqHD to detect correct antigen specificities using polyclonal CD8^+^ T cells sorted and stimulated with seven known antigens including potentially cross-reactive epitopes (**Supplementary Table 1**). PE-labeled, DNA-barcoded tetramers were used to stain cultured T cells. Tetramer^+^ CD8^+^ T cells were sorted (**Supplementary Fig. S1**) and loaded to BD Rhapsody^™^ to perform reverse transcription and PCRs. A total of 4,533 single cells were recovered after sequencing (**Supplementary Table 2**). Further filtering of low-quality cells and putative multiplets led to 4,462 cells retained, among which, a median of 140 genes were detected and TCRα and TCRβ capture efficiencies were 89% and 94% respectively. For each of these six polyclonal CD8^+^ T cell cultures, our previously developed MIDCIRS technology^20^ was used to assess TCRβ sequence diversity and distribution. These TCRβ sequences were then set as internal references for identifying true antigen specificities (**Supplementary Table 3**). Although the tetramer negative cells had a lower level of target gene expression, a similar level of gene expression were observed among different antigen-specific T cell clones (**Supplementary Fig. S2a, b**). An average of 17,249 reads per cell were sequenced for tetramer DNA-barcodes.

We detected antigen binding events based on MID count distribution of tetramer DNA-barcodes in each single cell, which helped us to define antigen specificity and possible cross-reactive binding antigens for individual T cells (**See methods**). Using the known TCR sequences from T cell clones, their known antigen specificities, and detected antigen-specificity by pMHC DNA-barcode, we showed that the precision, which is antigen-matched TCRs divided by antigen-specific TCRs identified by pMHC DNA-barcodes, is over 98% and the recall, which is antigen-matched TCRs divided by all TCRs determined by TCRβ clonality, is over 80%, except for GAD specific clones (**Fig 2a, b**). Additional analysis revealed the lower recall rate for GAD specific clones was due to one non-GAD binding clone (TCRβ: CASRFLGTEAFF) that accounted for 26% of all GAD specific T cells, which is likely to be a non-specific contaminant in the polyclonal culture (**Fig. 2c-h**).

**Figure 2.**
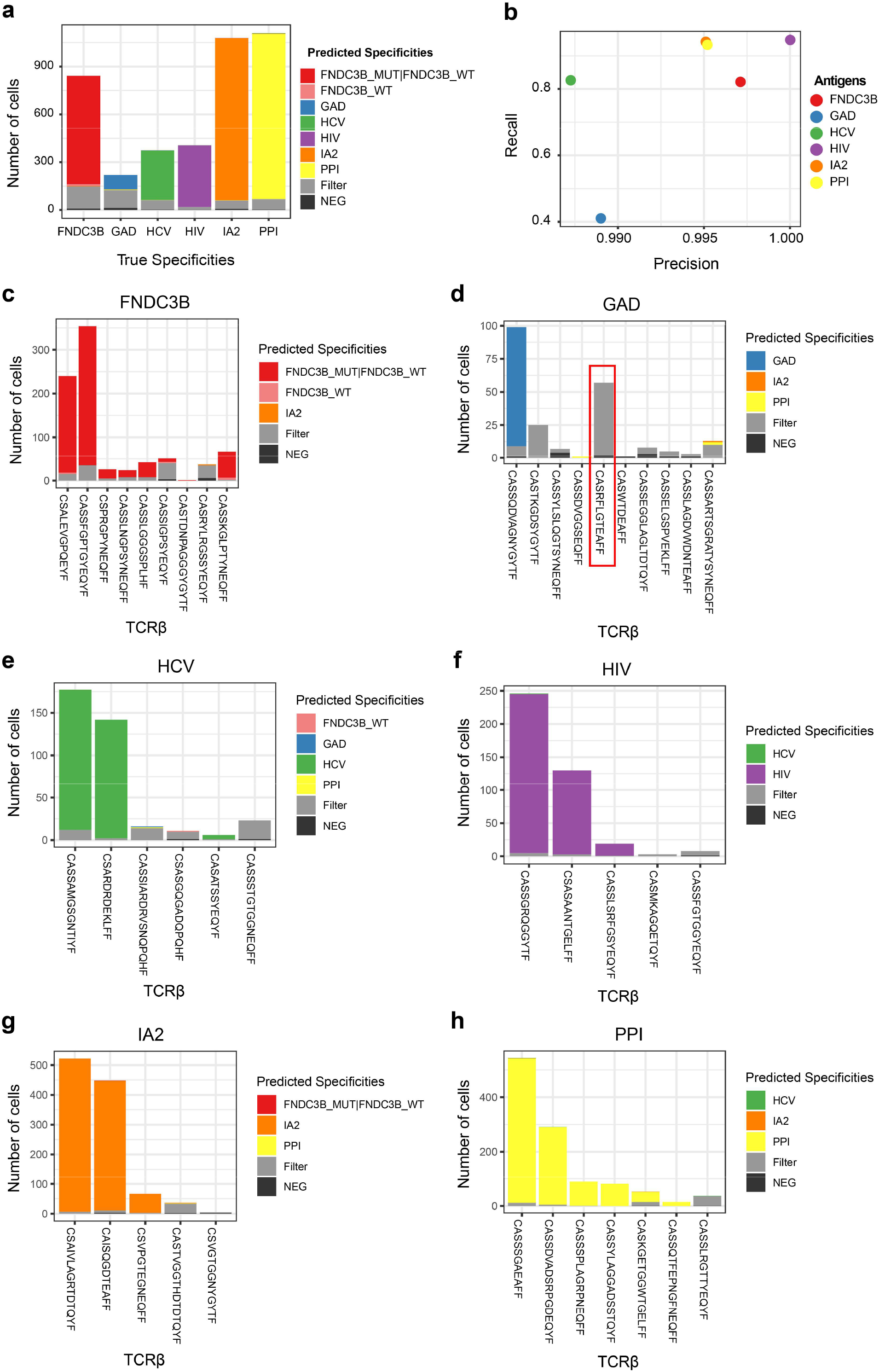
TetTCR-SeqHD validation in cultured polyclonal CD8^+^ T cells. Seven pMHCs were used to sort seven groups of polyclonal T cells and expanded *in vitro*. These included a group of cross-reactive T cells sorted by two similar antigens, FNDC3B_WT and FNDC3B_MUT. TCRβ sequencing was performed for each polyclonal T cell culture using MIDCIRS. These TCRs and their associate antigen specificity were used to assess the recall and precision rates (see **Methods**) for TetTCR-SeqHD. (**a**) Fraction of antigen specificities identified in different categories for each of the six polyclonal T cell cultures. True specificities was assigned based on TCRβ sequence found in the TetTCR-SeqHD experiment that matches known TCR sequences from bulk TCRβ sequencing. Cells were classified into the “filter” category based on the criteria described in **Methods**. (**b**) The recall and precision rates for each polyclonal T cell culture shown in (**a**). (**c**)-(**h**) Distribution of predicted antigen specificities for each T cell clone within each polyclonal T cell culture. The x-axis for each plot was ranked by TCRβ associated transcript copy numbers from MIDCIRS assay (Left to right, high to low). Red box denotes the contaminant TCR clone in GAD antigen-specific polyclonal culture. Filter category is defined same as in (**a**).

### Diverse T cell phenotypes revealed by TetTCR-SeqHD

To further demonstrate the advantages of TetTCR-SeqHD in characterizing antigen-specific CD8^+^ T cells, we curated a panel of 215 endogenous and 65 foreign antigens from the IEDB database and based on peptide-MHC class I binding prediction (**Supplementary Table 4, see methods**) covering HLA-A01:01, HLA-A02:01, and HLA-B08:01 alleles and applied TetTCR-SeqHD in ten non-type 1 diabetes (T1D) healthy donors and eight T1D patients (**Supplementary Table 5**). Endogenous and foreign peptides were UV-exchanged^18^ onto PE and APC-labeled tetramers respectively. CD8^+^ T cells were stained and sorted similar to the polyclonal T cell cultures (**Supplementary Fig. S3**). An HCV antigen-specific CD8^+^ T cell clone was spiked in to primary CD8^+^ T cells for all HLA-A02:01 donors. A total of 35,168 cells were recovered across four experiments. An average of 50,000 reads per cell were sequenced covering all six groups of attributes (**Supplementary Table 2**). After single cell quality filtering and removing putative multiplets (See Methods), 32,992 cells were retained with a median of 62 detected genes and 47 detected antibodies per cell. Among all primary cells, 45% and 68% cells had TCRα and TCRβ captured, with pairing efficiency of 34%. Since the primary CD8^+^ T cells were recovered from frozen samples, lower gene and TCR capture rates were seen compared with cultured clones.

We started by performing joint modeling of RNA expression and surface-protein expression using totalVI^21^, followed by dimensionality reduction using uniform manifold approximation and projection (UMAP)^22^ and single cell clustering with the Leiden algorithm (**Fig. 3a**)^23^. Minimum batch effects among chips were detected (**Fig. 3b**). A total of 13 clusters were identified, consisting of major conventional CD8^+^ T cell phenotypes including naïve T cells (T_naïve_, clusters 1-4), central memory T cells (T_cm_, cluster 6), effector memory T cells (T_em_, clusters 8-10), effector T cells (T_eff_, clusters 11-12) and transitional T cells between effector and memory populations (T_trans_, cluster 7) based on CCR7 and CD45RA/CD45RO protein expression, spike-in HCV specific clone (cluster 13) and CD56^+^ T cells, which are likely to be NK-like T cells^24^ (cluster 5) (**Fig. 3c, d**). The large number of primary CD8^+^ T cells processed and the combined analysis of target gene and surface-protein expression provided a superior resolution to identify sub-populations. While clusters 8, 9, and 11 represent early stages of T_em_ and T_eff_, clusters 10 and 12 represent late stage T_em_ and T_eff_ based on the graduate changes of gene/protein expression. Similarly, T_naive_ was also further separately into four clusters (1-4) (**Fig. 3d, Supplementary Fig. S4**). Of note, we found that cluster 5 (CD56^+^ T cells) is characterized by a low tetramer DNA-barcode signal fraction (**Supplementary Fig. S5**), and no enrichment of antigen-specific CD8^+^ T cells was identified. Among the 12 clusters of primary CD8^+^ T cells identified from all donors, the four T_naïve_ clusters and the CD56^+^ T cell cluster have the lowest TCR clonality, which is ubiquitous in all donors. However, different activated T cell subpopulations display various degrees of clonal expansion and clusters 8, 9, 10, and 12 (T_em_ and T_eff_) have a relatively high TCR clonality in majority of donors (**Fig 3e**).

**Figure 3.**
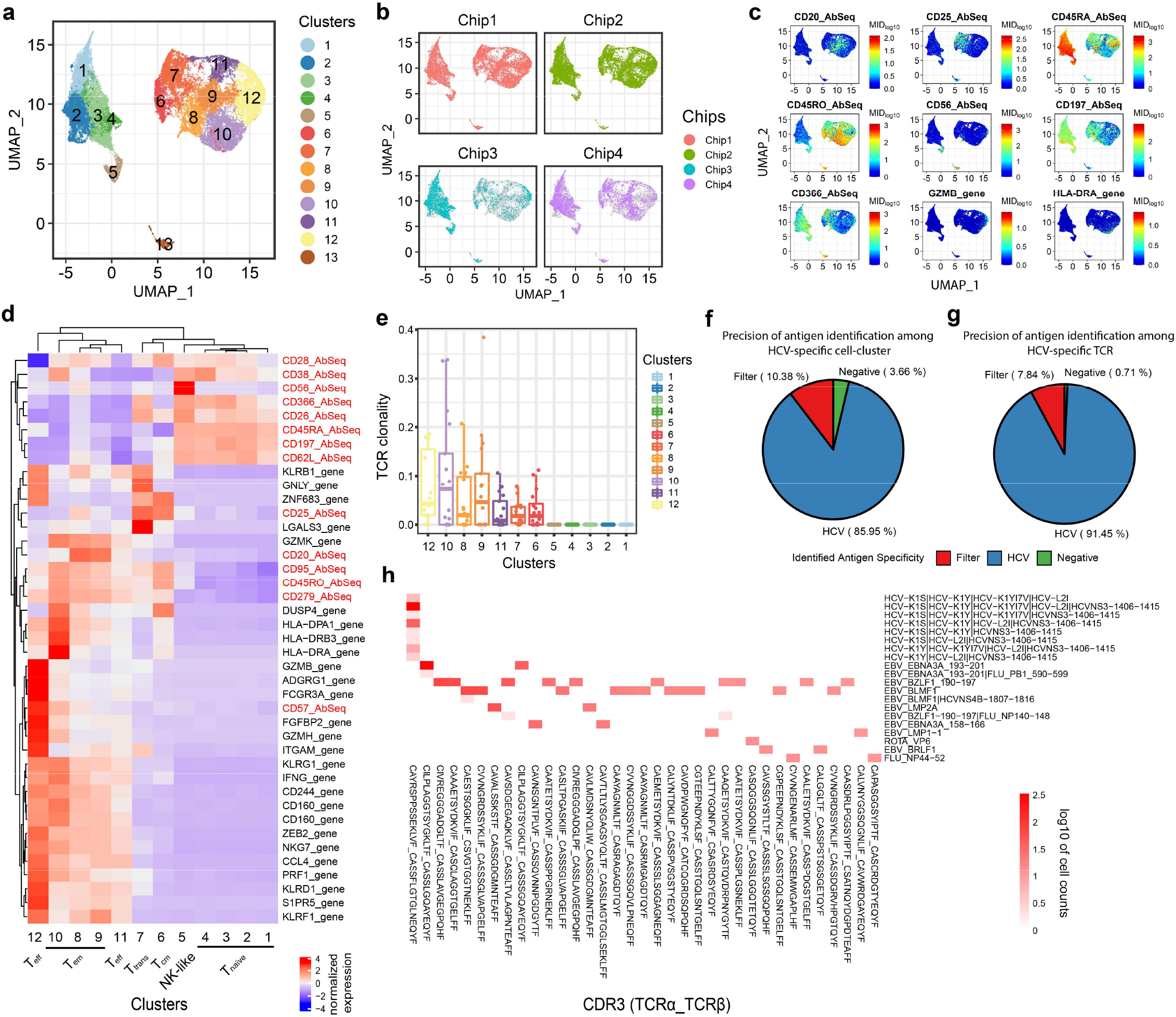
TetTCR-SeqHD enables combined gene expression, phenotype, and TCR clonality comparison among antigen-specific CD8^+^ T cells. (**a**) The UMAP projection of 32,992 single cells sorted from healthy and T1D donors. 13 clusters, including a cluster consisting of HCV spike-in T cell clone, were identified. (**b**) UMAP projection of single cells from different chips. Grey dots represent all cells and colored dots are cells from different chips. (**c**) Expression level of seven surface-proteins, CD20, CD25, CD45RA, CD45RO, CD56, CD197 (CCR7), and CD366 (TIM3), and two genes, *GZMB and HLA-DRA* across single T cells illustrated in (**a**). (**d**) Z-score normalized mean expression of differentially expressed genes and surface proteins (by antibodies) in each identified cluster. (**e**) TCR clonality in 12 primary CD8^+^ T cell clusters among 18 donors. (**f**)-(**g**) Precision of antigen identification among HCV-specific T cell-cluster (cluster13) (**f**) and HCV-specific TCR bearing cells (**g**). Cells were classified into “filter” category based on following criteria: 1) more than one antigens bind to single cell, and these antigens are more than 3 amino acid distance away from each other; 2) correlation of tetramer MID between single cell and median of all cells with same TCR sequence is below 0.9, identified as described in **Methods**. (**h**) Heatmap for the cognate antigen specificities of the top enriched TCRs (T cell clonality >= 10 cells). Top enriched TCRs are listed in the x-axis and the antigen specificities detected by TetTCR-SeqHD are listed in the y-axis. Colored blocks indicate antigen binding to a particular TCR. White background represents no binding event was detected for most of the TCR-antigen combinations.

### Foreign antigen-specific T cells display distinct phenotypes and degree of clonal expansion

Different donors show distinguishable phenotypic distributions on the UMAP projection (**Supplementary Fig. S6**), which prompted us to further examine the heterogenous functional profiles of antigen-specific T cells among donors. Altogether, 12,518 viral antigen-specific, 3,626 non-T1D-related endogenous antigen-specific and 1,952 T1D-related endogenous antigen-specific T cells were detected but the ratio varies in different individuals (**Supplementary Table 6**). Almost all of the clonally expanded TCRs had unique antigen specificities identified, confirming the precision of TetTCR-SeqHD in primary CD8^+^ T cells from human PBMCs (**Fig. 3f**). We further used the HCV clone to characterize the precision and recall of TetTCR-SeqHD in primary CD8^+^ T cell experiments. Of the cluster 13 identified to harbor the HCV specific spike-in clone, there were total of 623 cells, 536 (86%) of which were accurately identified as binding to at least one HCV wildtype (WT) and associated variant antigens (**Fig. 3g**). Of these cells, a total of 421 cells were identified to have the same paired TCRα/β sequences as the HCV specific clone in this experiment. 91% of them bind to at least one HCV wildtype (WT) and associated variant antigens (**Fig. 3h**). 40 viral antigens that were detected in greater than 5 cells across all donors were selected for further analysis (see **Methods**). As expected, different T cell phenotypic clusters are comprised of distinct antigen specificities, with endogenous antigens occupying T_naïve_, while foreign antigens populating non-naïve T cell clusters (**Fig. 4a**). In general, different donors, regardless of their T1D status, presented varying frequency and phenotypic profiles of viral antigen-specific CD8^+^ T cells, possibly due to different infection or vaccination history (**Fig. 4b-e**). However, we also found that some viral antigens induced distinct T cell phenotypes. Influenza antigen experienced T cells are mostly within cluster 7, where T cells display Tim3^+^, CD25^+^ and CD26^+^ phenotype^25-27^ (**Fig. 4c, Supplementary Fig. S7**). Epstein-Barr Virus (EBV) antigens showed distinguishable phenotypes compared with influenza antigens (**Fig. 4c, Supplementary Fig. S8**). Two different categories of EBV antigens originated from lytic and latent viral proteins also present distinct phenotypes. Antigens from latent viral proteins, such as LMP1 and LMP2, preferentially induced T cells in central memory states (cluster 6), while lytic viral proteins, such as BRLF1 and BLMF1, display effector and effector memory phenotypes (clusters 8, 9, 10, and 12) (**Fig. 4c**), consistent with previous findings using CyTOF^16^. We also found M1 specific CD8^+^ T cells display a more uniform phenotype distribution among donors, compared with other antigens (**Fig. 4d, e**).

**Figure 4.**
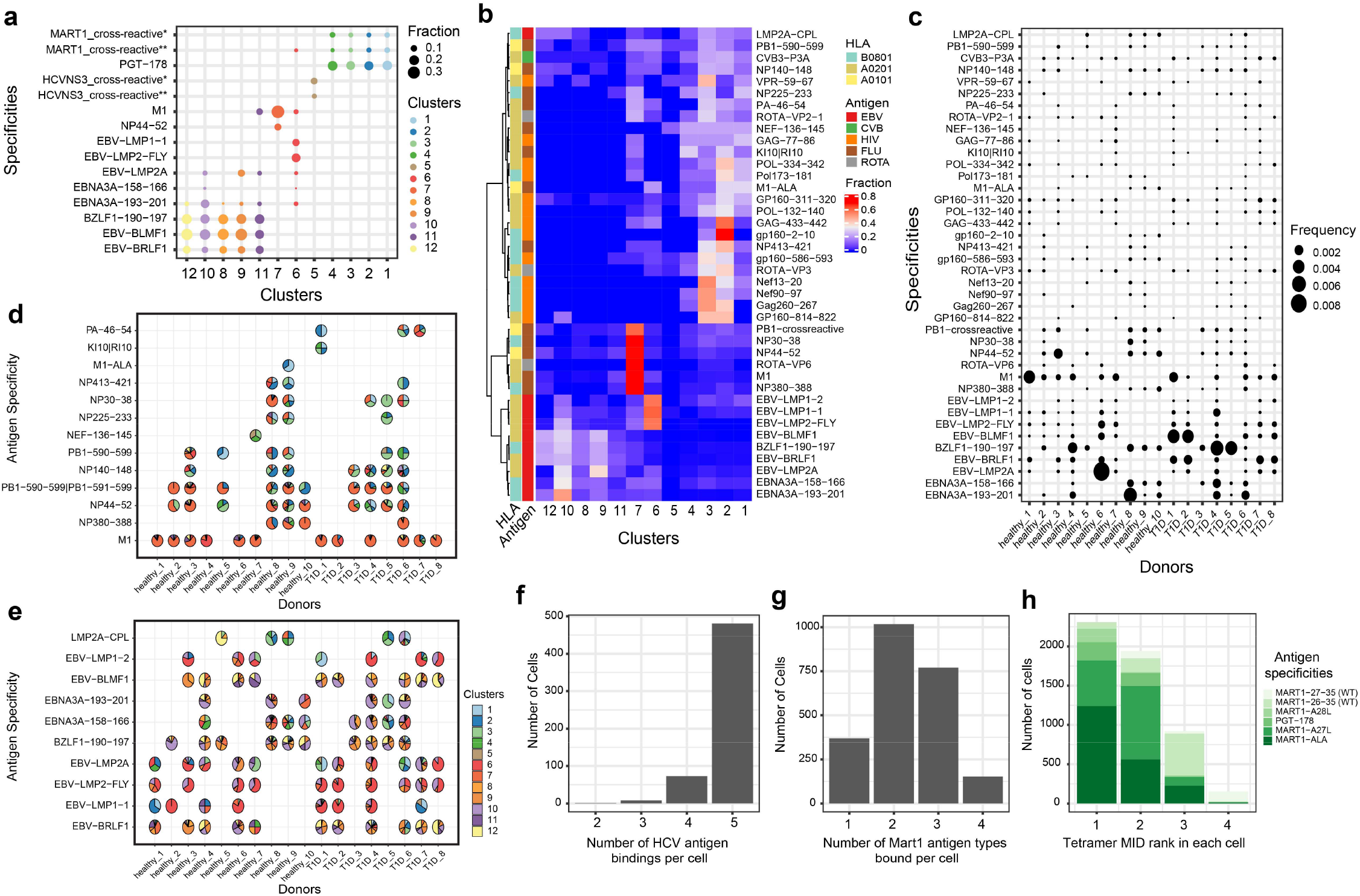
Gene expression and phenotype analysis of foreign-antigen specific T cells and cross-reactivity validation. (**a**) The top representative antigen specificities in 12 primary CD8^+^ T cell clusters (MART1_crossreactive*: MART1-26-35|MART1-A27L|MART1-ALA; MART1_crossreactive**: MART1-A27L|MART1-ALA; HCVNS3_crossreactive*: HCV-K1S|HCV-K1Y|HCV-K1YI7V|HCV-L2I|HCVNS3-1406-1415; HCVNS3_crossreactive**: HCV-K1S|HCV-K1Y|HCV-L2I|HCVNS3-1406-1415). (**b**) The distribution of viral-antigen specific CD8^+^ T cells from all 18 donors among 12 primary CD8^+^ T cells clusters. (**c**) Frequency comparison of viral-antigen specific T cells in 18 donors (in **b**,**c**: PB1-crossreactive represents cross-reactivity between PB1-590-599 and PB1-591-599). (**d**)-(**e**) Phenotypes distribution of influenza-specific (**d**) and EBV-specific (**e**) T cells in each individual. Each pie represents the T cell distribution in 12 phenotypes of primary CD8^+^ T cells for the corresponding donor (x-axis) and antigen (y-axis) combination. Empty spaces **d** and **e** mean no antigens were detected for the corresponding donor-antigen combination. (**f**) Histogram of the number of different types of HCV antigens, WT and variant antigens, bound per cell. (**g**) Histogram showing the number of different types of Mart1 antigens, WTs and variant antigens, bound per cell. (**h**) Distributions of tetramer MID ranks of Mart1 WT, variant antigens, and a cross-reactive antigen in each cell for four groups of binding patterns. MART1-27-35: AAGIGILTV (WT), MART1-26-35: EAAGIGILTV (WT), MART1-A28L: ALGIGILTV, MART1-A27L: ELAGIGILTV, MART1-ALA: ALAGIGILTV, PGT-178: LLAGIGTVPI, PGT-178: LLAGIGTVPI.

Another advantage of TetTCR-SeqHD is its capacity to identify putative cross-reactive CD8^+^ T cells. Similar to TetTCR-Seq^18^, 85% of HCV specific clone display binding to all five HCV antigens^18^ (**Fig. 4f**). We also examined the cross-reactivity detection in primary CD8^+^ T cells using MART1 antigens. MART1 wildtype antigens (MART1_27-35_ nonamer and MART1_26-35_ decamer) and its variants have been widely used as a model system of human cancer antigens. By changing one or two amino acids, such as MART1_26-35_ A27L and MART1_26-35_ E26A/A27L, it was noted previously that the resulting variant peptides greatly improved the binding/stability of peptide/HLA-A*0201 complexes and enabled the otherwise weak wildtype antigens to potent immunogens^28, 29^. We thus used these set of peptides and studied the robustness of TetTCR-SeqHD in detecting both strong and weak pMHC ligands. Among cells with same tetramer binding frequency greater than 10, a total of 2,308 cells were identified to bind MART1 WT or variant antigens. 84% of cells bind to more than one MART1 WT or variant antigens (**Fig. 4g**). Interestingly, our method also detected previously noted cross-reactivities among the PGT-178 (LLAGIGTVPI) peptides and a MART1 variant antigen (ELAGIGILTV)^30^ and an additionalMART1 variant cross-reactive antigen (ALAGIGILTV), despite five or more amino acid differences in these peptides (**Fig. 4h**).

### Selected autoantigens exhibit differences between healthy and T1D patients

Among 209 T1D-related autoantigens included in the antigen pool, 106 and 102 different autoantigens were detected more than 3 times in 1,109 and 814 T1D antigen-specific cells from T1D and non-T1D donors, respectively. The total T1D autoantigen tetramer^+^ CD8 T cell frequency was comparable between T1D and healthy donors (**Supplementary Fig. S9**). However, comparing the frequency of T1D autoantigen-specific CD8^+^ T cells individually, we found INS-WMR-10, PPI-29-38 and PTPRN-805-813 specific cells exhibit a significantly higher cell frequency in T1D patients compared to healthy control donors within this donor cohort (**Fig. 5a**). Among them, PTPRN-805-813 was reported before as a potential marker in PBMC of T1D patients^31^ and PPI-29-38 was identified as an HLA-A02:01 low binder but present in T1D patients^32^. To ensure the sensitivity of our analysis, we increased the tetramer MID negative threshold to 15 and compared the frequency difference between T1D and healthy donors again. Five antigens were identified, including previously identified INS-WMR-10 and PTPRN-805-813, further validating the potential of these two antigens to distinguish between T1D and healthy donors (**Supplementary Fig. S10a**). We also noticed varying degree of clonal expansion in T1D autoantigen-specific T cells isolated from different T1D patients, revealing the complexity of antigen landscape in T1D (**Supplementary Fig. S10b**). This could also be caused by limited sampling from PBMC.

**Figure 5.**
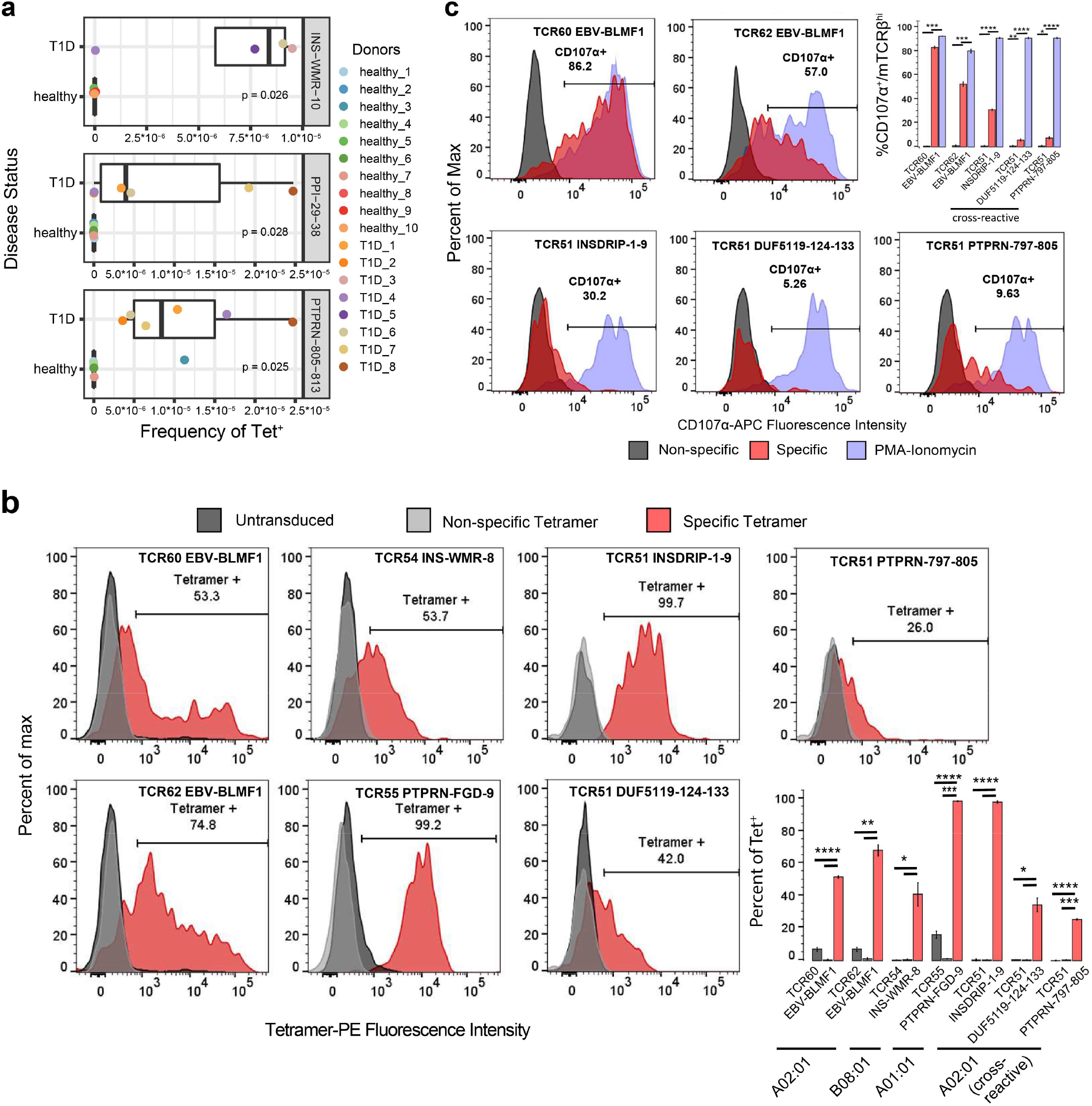
Identification of three T cell specificities selectively enriched in T1D patients and TCR specificity and cross-reactivity validation. (**a**) Three T1D-related antigens (INS-WMR-10, PPI-29-38, and PTPRN-805-813) were identified to have a significantly higher frequency of cognate T cells in the peripheral blood of T1D patients compared to healthy donors. Wilcoxon signed-rank test was performed. (**b**) TCR specificity and cross-reactivity validation by tetramer staining. Five TCRs that were identified to recognize six different antigens in complex with distinct HLA alleles, including TCR51 that recognized three unrelated antigens, were transduced into human primary CD8^+^ T cells and stained with respective tetramers. HLA-A02:01: EBV-BLMF1, INSDRIP-1-9, DUF5119-124-133, PTPRN-797-805; HLA-B08:01: INS-WMR-8; HLA-A01:01: PTPRN-FGD-9. (**c**) TCR specificity and cross-reactivity validation by T cell functionality. The HLA-A02:01 restricted TCR transduced cells generated in (**b**) were further stimulated with their cognate antigens and the expression of CD107α was measured on TCRβ^hi^ fraction of the cells. Two-tailed student’s *t*-test was performed. EBV-BLMF1: GLCTLVAML; INSDRIP-1-9: MLYQHLLPL; DUF5119-124-133: MVWGPDPLYV; PTPRN-797-805: MVWESGCTV; INS-WMR-8: WMRLLPLL; PTPRN-FGD-9: FGDHPGHSY. *, p ≤ 0.05; **, p ≤ 0.01; ***, p ≤ 0.001; ****, p ≤ 0.0001.

In addition, we identified an expanded T cell clone cross-recognizing three different antigens, INSDRIP-1-9, DUF5119-124-133 and PTPRN-797-805 in a type 2 diabetes patient. This led us to test the plasma banked from the same blood draw and showed the patient was positive for GAD (Glutamic Acid Decarboxylase) reactive auto-antibody. Further review of the medical record showed that the patient was later diagnosed with Latent Autoimmune Diabetes in Adults (LADA) after the sample was collected for this study. Interestingly, INSDRIP-1-9 is derived from an alternative open reading frame within human insulin mRNA and a significantly higher levels of INSDRIP-1-9^+^ specific CD8^+^ T cells were reported to be detected in T1D patients ^33^. DUF5119-124-133 is derived from *Bacteroides fragilis /thetaiotaomicron*, a common bacteria found in human gut microbiota^34^ and PTPRN-797-805 is derived from IA2 protein, a previously known T1D autoantigen^35^. This is likely due to cross-reactivity of the three antigens by the same TCR. To confirm the result by TetTCR-SeqHD, we thus transduced this TCR together with some TCRs identified among T1D and healthy subjects to further validate the accuracy of TetTCR-SeqHD (**Supplementary Table 7**). Tetramer staining (**Fig. 5b**) and antigen stimulation experiments (**Fig. 5c**) both confirmed that cognate TCRs identified from TetTCR-SeqHD can bind and be stimulated by respective antigens.

## Discussion

In this study, we developed a method to simultaneously profile TCR sequences, cognate antigen specificity, gene expression, and surface-protein expression, for single primary CD8^+^ T cells in a high throughput manner. We addressed the precision of TetTCR-SeqHD, ability to profile TCR cross-reactivity, as well as its application to study diverse phenotypes of foreign- and self-antigen specific CD8^+^ T cells. By using *in vitro* cultured polyclonal T cells with known antigen specificities and TCR sequences, TetTCR-SeqHD established over 98% precision for detecting the correct antigen specificity.

Recently, DNA-barcoded dextramer, dCODE^™^ dextramer^®^ was adapted by 10x Genomic platform to enable profiling of antigen-specific CD8+ T cells. However, the dCODE^™^ dextramer^®^ suffers from the high cost of generation of dextramers, thus lacks the flexibility to screen large antigen panels. This prevents it from profiling antigens in a high-throughput manner. By combining *in vitro* transcription and translation (IVTT) with UV exchange technique, TetTCR-SeqHD enables the creation of a panel of antigens (in hundreds) affordably and quickly (within a week). Therefore, we created a large panel of antigens consisting of foreign-specific antigens derived from various virus and self-specific antigens derived from known T1D autoantigens, and profiled CD8^+^ T cells recognize these antigens from healthy subjects and T1D patients.

With the ability to profile targeted gene expression and surface-protein expression simultaneously using BD Rhapsody^™^ platform, we resolved 12 clusters for primary CD8^+^ T cells, plus 1 cluster for *in vitro* cultured HCV-specific CD8^+^ T cell clone. Most importantly, T cell phenotypic and functional sub-classes represented by gradual changes of gene expression were revealed among these 12 clusters, from naïve to early stage of effector and memory populations to transitional state between effector and memory, and to late stage of effector and memory populations. By investigating the composition of phenotypic clusters for each antigen, phenotype signatures of distinct antigens were assessed. We found that viral antigens from influenza and EBV display distinct phenotypes. Influenza-specific CD8^+^ T cells were mostly enriched in cluster 7, displaying a transitional phenotype between effector and memory populations, while EBV-specific CD8^+^ T cells were largely memory and effector populations. Similar phenotypic differences between EBV latent and lytic antigens were observed previously using mass cytometry^16^. This example further validates the robustness of TetTCR-SeqHD to capture the phenotypic profiles of antigen-specific CD8^+^ T cells. Moreover, studied subjects also showed diverse phenotype signatures of influenza- and EBV-specific CD8^+^ T cells, due to different viral infection (or vaccination) history.

In addition to its high precision and high-throughput capacity, TetTCR-SeqHD also enables detection of cross-reactivity of CD8^+^ T cells. We examined cross-reactivity in both *in vitro* cultured HCV-specific CD8^+^ T cell clones and primary CD8^+^ T cells. We not only detected cross-reactivity among HCV and MART1 WT and variant antigens, but also found cross-reactivity among INSDRIP-1-9, DUF5119-124-133 and PTPRN-797-805 in a type 2 diabetes patient. The TCR sequences obtained simultaneously demonstrated their critical role in validate antigen-specificity and cross-reactivity in high-throughput antigen screening and antigen-specific T cell profiling. Interestingly, these three antigens are more than 3 amino acid away from each other, underscoring the flexibility of TCR-antigen recognition between dissimilar peptides. Given that DUF5119-124-133 is derived from human gut microbiota, the association between certain dysbiosis of gut microbiome and the role of T cells in the onset of T1D requires further investigation.

Lastly, with the panel of T1D autoantigens, we investigated the differences of autoantigen-specific CD8^+^ T cells between healthy and T1D subjects. Although, we did not identify any phenotypic differences (data not shown), we found three antigens, INS-WMR-10, PPI-29-38 and PTPRN-805-813 which exhibit a significantly higher antigen-specific CD8^+^ T cell frequency in T1D patients within this donor cohort. Wiedeman et al. recently found that activated islet-specific CD8^+^ memory T cells were prevalent in subjects with T1D who experienced rapid loss of C-peptide while slow disease progression was associated with an exhaustion-like profile^36^. In contrast, Culina et al. reported predominant naïve phenotype for circulating islet-specific CD8^+^ T cells in T1D^37^, similar to our results. These contradicting results are likely due to different patient cohorts with different T1D onset timing as well as choice of T1D antigens. Given that a similar attempt using a much smaller panel of T1D related auto-antigens failed to identify any antigens within PBMCs that would separate healthy individuals from T1D patients ^37^, our results provide an interesting premise that warrants further tests in a much larger cohort, which could be very useful in T1D early diagnosis.

Due to the advantage of multi-dimensional profiling of single cells, TetTCR-SeqHD method enables one to identify phenotypic differences of antigen-specific CD8^+^ T cells, distinguish disease status, screen antigens with a high-throughput, and identify TCRs with therapeutic potential. TetTCR-SeqHD is likely be a game changer in basic and translational research focusing on T cells.

### Methods

#### 1. Generation of DNA-barcoded fluorescent streptavidin

The conjugation of DNA linker (**Supplementary Table 8**) to PE- or APC-labeled streptavidin was performed as previously described with slight modifications^18^. During S-HyNic modification of PE- or APC-labeled streptavidin, 2 moles equivalent of S-HyNic were used. Following the conjugation of DNA linker, peptide-encoding DNA-barcodes (**Supplementary Table 4**) were annealed to the complementary DNA linker on the DNA linker PE or APC streptavidin conjugate in the presence of 1x NEBbuffer2 (NEB) with below programs: 60°C for 30s, then -1°C/cycle for 35 cycles. The final DNA-barcoded fluorescent streptavidin conjugate was stored at 4°C.

#### 2. In vitro transcription and translation

Peptide-encoding DNA oligonucleotides were purchased from Sigma Aldrich. 50nM DNA templates were first amplified by PCR as described previously with modifications (Zhang et al.). 1µM IVTT_r and IVTT_f primers (**Supplementary Table 8**) were used following below reaction conditions: 95°C 3min; then 22 cycles of 95°C for 20s, 59°C for 30s, 72°C for 30s; then 72°C for 5min. The PCR product was then diluted with 50µl of nuclease free water and proceeded to IVTT reaction.

#### 3. Generation of pMHC tetramer library

IVTT generated peptides were mixed with biotinylated pMHC containing a UV-labile peptide. The final concentration of biotinylated pMHC is 0.2mg/ml. Individual pMHC was formed through UV exchange as described previously^38^. Individual pMHC tetramer and tetramer library pool were generated and tested as described previously^18^. Tetramer pool can be stored in 4°C temporarily.

#### 4. Customization of CD2 SampleTag, custom AbSeq, and custom CD50 SampleTag

Anti-CD2 antibody was purchased from Biolegend (Clone RPA-2.10, Biolegend). Amine modified oligonucleotide was purchased from Sigma Aldrich (**Supplementary Table 8**). The conjugation between oligonucleotide and CD2 antibody followed CITE-Seq protocol^4^.

Corresponding antibodies and used oligonucleotides were listed in supplementary table (**Supplementary Table 9**).

12 CD50 antibody SampleTags^39^ were customized by BD Biosciences using the commercial SampleTag oligos.

#### 5. Sorting and culture of antigen-specific CD8^+^ T cell polyclones

Seven types of tetramers with peptides chemically synthesized and UV-exchanged to MHC were used to raise antigen-specific polyclonal T cells (**Supplementary Table 1**). For each tetramer, 20 Tetramer^+^ CD8^+^ single T cells were sorted into each well of the 96 well plate for culture for three weeks.

#### 6. pMHC tetramer staining and sorting of primary human CD8^+^ T cells

Human whole blood from diagnosed T1D and T2D patients were obtained at Seton Family of Hospitals at Austin with informed consent. The use of whole blood from these patients was approved by the Institutional Review Board of the Ascension Seton University Physicians Group and is compliant with all relevant ethical regulations. Human peripheral blood mononuclear cell (PBMC) from healthy donors were purchased from ePBMC. PBMC from T1D whole blood was isolated using Ficoll-paque density-gradient centrifugation (GE Healthcare). CD8^+^ T cells were then enriched from PBMC of T1DM and healthy donors using EasySep™ Human CD8^+^ T cell isolation kit (STEMCELL).

CD8^+^ T cells were resuspended in FACS buffer containing 0.05% sodium azide and 50nM of Dasatinib. CD8^+^ T cells were then incubated at 37°C for 30min-60min. About 10,000 cells from an HCV peptide binding clone used previously^18^ were pre-stained with BV510 anti-CD8a antibody (clone: RPA-T8, Biolegend) and spiked into the primary CD8^+^ T cells. Following the Dasatinib treatment, tetramer pool together with anti-CD8a antibody (clone: RPA-T8, Biolegend) was directly added into the cells. Cells were incubated at 4°C for 1hr with continuous rotation. After washing, cells were further stained at 4°C for 20min with the presence of 5 µg/ml mouse anti-PE (clone: PE001, Biolegend) and/or mouse anti-APC (clone: APC003, Biolegend). AbSeq staining mastermix was prepared by pooling 1µl of each AbSeq together (**Supplementary Table 9**). Cells were washed in FACS buffer once and stained with the AbSeq mastermix. Additional dump-channel antibodies (AF488-anti-CD4, AF488-anti-CD14 and AF488-anti-CD19), 7-AAD and 2µl of anti-CD50 SampleTag were mixed in cells. Cells were incubated at 4°C for 40mins, prior to washing in FACS buffer twice and proceeded for sorting.

During cell sorting, about 50,000 Tet^-^CD8^+^ T cells were also sorted, which will later be spiked into Tet^+^ T cells.

#### 7. BD Rhapsody^™^ sequencing library preparation and sequencing

Prior to BD Rhapsody^™^ processing, Tetramer^-^CD8^+^ T cells were first stained with 2µl of CD2 SampleTag at 4°C for 30mins. Cells were washed in FACS buffer for three times and resuspended in 100µl BD Sample Buffer. Sorted Tet^+^CD8^+^ T cells and Tetramer^-^CD8^+^ T cells were counted using BD Rhapsody^™^. Tetramer^+^ and Tetramer^-^ CD8^+^ T cells were pooled and processed on BD Rhapsody^™^ cartridge following user’s manual. Single cell cDNA synthesis and library amplification were performed following manufacturer’s protocol with some modifications. Briefly, in PCR1, 1.2µl of tetramer PCR1 primer was added to the PCR reaction in addition to primers for gene expression panel, AbSeq, SampleTag, and universal oligo (**Supplementary Table 9**). 9 and 10 PCR cycles were used for 5000-10,000 and 10,001-20,000 cells respectively. Double-sided AMPure beads purification was processed to purify short amplicons (AbSeq, SampleTag and tetramer DNA-barcodes) and long amplicons (target genes and TCRα/β) separately. In PCR2, five separate PCR reactions with 15 reaction cycles were carried out to amplify gene panel, SampleTag, TCRα, TCRβ, and tetramer DNA-barcodes. AbSeq, tetramer and TCRα/β libraries were gel extracted for the desired band before proceeding to PCR3. Finally, 8 cycles of PCR reactions were performed for all six elements following manufacturer’s instruction. All PCR libraries were quantified using Bioanalyzer 2100 and pooled. 15% PhiX was used in all sequencing runs. Pooled libraries were sequenced on HiSeq X with PE150.

#### 8. BD Rhapsody^™^ sequencing pre-processing

Sequencing reads from target gene expression, AbSeq, SampleTag, TCRα/β and tetramer DNA-barcodes were processed as below (**Supplementary Fig. S11**).

For target gene expression and AbSeq sequencing, reads were processed with *BD Targeted Multiplex Rhapsody Analysis Pipeline Version 1*.*5* on Seven Bridges platform following manufacturer’s instructions. True cell barcodes were converted to oligonucleotide sequences according to BD cell barcode indexing rule. Then sequencing data of Tetramer, TCRα and TCRβ were processed using umitools^40^ to extract cellular barcode and unique molecular identifier (MID) for each read. Reads that are mapped to true cell barcodes were obtained.

For tetramer DNA-barcodes, only reads that are exact match of tetramer DNA-barcode reference were retained. Number of reads of the same MID tagged tetramer DNA-barcode (unique tetramer DNA-barcode) were counted for each cell. The distribution of the reads of unique tetramer DNA-barcode follows a bimodal distribution as reported previously^18^. The first peak corresponds to PCR and sequencing errors, and thus, reads falling under the first peak were filtered. Further the number of MIDs aligned to each tetramer DNA-barcode in each cell were counted to construct tetramer DNA-barcode count matrix.

For the SampleTag DNA-barcodes, reads were mapped to SampleTag DNA-barcode reference using bowtie2 with --norc and --local mode^41^. Aligned reads were then processed using umi_tools to count the number of MIDs for each SampleTag DNA-barcode in each cell. Distribution of MID counts for each SampleTag was fitted by a bimodal distribution and the cutoff between two distributions were set as the negative threshold for the corresponding SampleTag. In addition, to recover false negative SampleTag signals, SampleTag, whose MID counts account for >50% total SampleTag MID counts, was also classified as positive event. Cells containing CD2 SampleTag were Tet^-^ cells, while cells with more than two regular SampleTags were multiplets and were removed from further analysis.

For the TCR sequencing reads, we adapted sub-clustering algorithm as previously described^42^to remove PCR and/or sequencing errors and identify VDJ and CDR3 with some changes. Reads were first aligned to TCR J and C region reference. Only reads that are >62.5% identical were retained. Reads with same cellular barcodes and MID were grouped together. Under each group, reads within a Levenshtein distance of 15% were further clustered into a subgroup. For each subgroup, a consensus sequence was built based on the average nucleotide at each position, weighted by quality score. After ranking the consensus sequences by their abundance, the most abundant consensus sequence is selected and other sequences with edited distance less than three were removed. In case the most abundant consensus sequence is non-productive, the next most abundant productive sequence, if exists, was selected as the unique consensus sequence for that cell. The 2^nd^ TCR chain was retained when its MID count accounts for more than 20% of total TCRα or TCRβ MID counts.

#### 9. Dimensionality reduction, clustering and differential expression of single cells

All single cells were first filtered to exclude low quality cells whose total gene and AbSeq expression MID counts were in the last 1% quantile. Then cells identified as multiplets with SampleTag and cells with two productive TCRβ chains were also removed. Additionally, genes or AbSeqs whose expression were detected in less than 50 cells were filtered. Gene expression and AbSeq data from different Rhapsody chips were pooled together and performed joint probabilistic modeling of RNA expression and surface protein measurement with totalVI^21^. Each donor was treated as an independent batch factor and 200 epochs were used to train the model. Other parameters were set as default in totalVI. Posterior dataset was then used for dimensionality reduction (UMAP algorithm) and clustering (Leiden algorithm), both with Scanpy^43^.

#### 10. Calling tetramer specificity for each single cell

First, for each tetramer fluorescent color, distribution of total tetramer DNA-barcode counts per cell was fitted to a bimodal distribution. The cutoff counts were set as negative threshold to capture positive tetramer binding events. Tetramer DNA-barcode counts were then ranked for each single cell and the knee point on the count-rank plot was selected. Antigens rank higher than the inflection point are putative binding antigens. Besides, antigens that rank below inflection point, but with <=3 amino acid difference compared with higher ranking antigens, were also included as putative cross-reactive binding antigens. For each cell, tetramer MID signal fraction was defined as the fraction of cumulative MID count from putative binding antigens over cumulative MID count from all bound antigens:

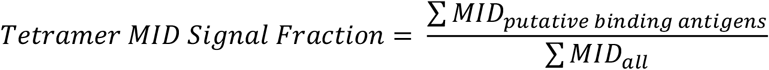

Further, cells with the same TCRα/β were pooled together. The correlation coefficient of antigen binding for each single cell in the pool were calculated between detected tetramer DNA-barcode counts and corresponding median tetramer DNA-barcode counts within the pool. This correlation coefficient for each single cell is used as the tetramer binding noise. The knee point of the distribution of correlation coefficients were set as the threshold, below which cells were removed due to high tetramer binding noise.

For analysis of viral antigens, we select antigens detected in more than 5 cells to ensure capturing low frequent antigen specific CD8^+^ T cells while limiting non-specific binding.

For sensitivity analysis to demonstrate the robustness of TetTCR-SeqHD, we set the negative threshold of tetramer MID to 15 to capture positive binding events. This threshold was then used for all experiments (**Supplementary Fig. S19**).

#### 11. Precision and recall rate calculation for TetTCR-SeqHD

In the TetTCR-SeqHD clone experiment, true positive is defined as antigen-matched TCRs between MIDCIRS and TetTCR-SeqHD. Predicted condition positive is defined as antigen-specific TCRs identified by pMHC DNA-barcodes. The condition positive is defined as antigen-specific TCRs identified by MIDCIRS. Precision and Recall is then calculated as below.

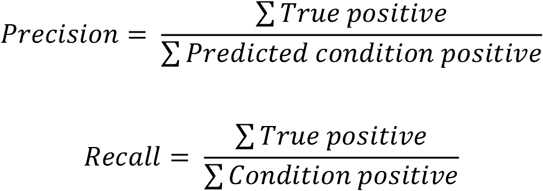

#### 12. Prediction of peptide-MHC class I binding

HLA-A02:01 bound T1D autoantigens were curated from the IEDB (www.iedb.org) database, while HLA-A01:01 and HLA-B08:01 bound T1D autoantigens were predicted using NetMHCpan 4.0^44^. The IC50 cutoff for HLA-A01:01 and HLA-B08:01 was 950nM and 500nM respectively.

#### 13. TCR clonality calculation

TCRs that have productive paired α and β chains were used to calculate TCR clonality, which is a score to characterize T cell expansion. Higher TCR clonality indicates that corresponding TCR are more clonally expanded. If there is singleton TCR, we define the TCR clonality being 0, while single TCR species with multiple copies have TCR clonality being 1. For all other situations, the TCR clonality is defined using following formula.

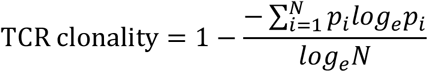

#### 14. TCR transduction

We generated TCR constructs as previously described^18^ and cloned them into an empty pCDH (System Biosciences) vector driven by the MSCV promoter. Lentivirus was generated using the Virapower (ThermoFisher Scientific) system and concentrated 10 times using an Amicon Ultra column. Freshly thawed CD8^+^ T cells from an HLA-A2/B8/A1 negative donor were stimulated with Immunocult (StemCell Technologies) and incubated with the concentrated virus for 2-3 days. The cells were expanded for a minimum of ten days and then assessed for murine TCRβ chain expression.

#### 15. Flow Cytometry on transduced cell

Tetramer staining was performed as previously described^18^ with tetrameric-MHC loaded with chemically synthesized peptides (Genscript). Briefly, the transduced cells and negative controls were stained with an anti-CD8a antibody (clone RPA-T8, Biolegend) before addition of tetramer for one hour on ice. Negative controls were established using non-specific tetramer (HLA-A*02:01:HCVns3:1406-1415 – KLVALGINAV) and un-transduced T cells from the same donor. After washing, the cells were stained with an anti-murine TCRβ antibody (Biolegend) and 7-AAD before analysis on a BD Accuri.

T2 cells were pulsed with chemically synthesized peptide (10µM) for 2 hours at 37°C. The cells were then washed and incubated 1:1 with the transduced cells for four hours at 37°C. Negative and positive controls were performed using non-specific peptide (HCVns3:1406-1415) and PMA/ionomycin (Cell Stimulation Cocktail, Biolegend), respectively. During incubation, anti-CD107α (Biolegend) antibody and monensin were added to detect and stabilize degranulation events. The assay was stopped via addition of cold PBS and subsequent staining for CD107α, CD8α, and murine TCRβ (Biolegend). Cells were analyzed via a BD Accuri.

#### 16. Detection of Auto-antibodies

The presence of anti-GAD, -IA2 and -Znt8 antibodies were determined via ELISA assay obtained from Kronus and performed according to the manufacturer’s instructions. Whole, undiluted plasma was used in this assay. Absorbance was measured using a SpectraMax M3 plate reader and analysis of the standard curve was performed in R using a cubic-spline fit. The antibody concentration for each sample was then interpolated, with all positive controls falling within the reported concentrations. Patients were reported as positive if the detectable antibody levels were in excess of 5 IU/mL, 7.5 and 15 U/mL for the anti-GAD, -IA2 and -Znt8 antibodies, respectively according to the manufacturer’s instructions.

## Acknowledgements

We thank T1D patients for donating blood samples to our study. We also thank anonymous blood donors and staff members at We Are Blood for sample collection. We thank P. Parker for assistance with blood sample purification. This work was supported by NIH grants S10OD020072 (N.J.) and R33CA225539 (N.J.), by NSF CAREER Award 1653866 (N.J.), by Welch Foundation grant F1785 (N.J.), by the Robert J. Kleberg, Jr. and Helen C. Kleberg Foundation (N.J.), and by the Chan Zuckerberg Initiative Neurodegeneration Challenge Network Ben Barres Early Career Acceleration Awards 191856 (N.J.). We would also like to acknowledge funding from the Cockrell School of Engineering Fellowship (A.S.), Mario E. Ramirez Endowed Graduate Fellowship (A.S.), and the Harry and Rubye Gaston Graduate Scholarship (A.S.).

## Author contributions

K.-Y.M. and N.J. conceived and designed the study. K.-Y.M. designed and developed the technology platform; K.-Y.M. and A.A.S. performed and analyzed data for the majority of experiments; K.-Y.M. developed pipeline to analyze tetramer DNA-barcode data; C.H. and K.-Y.M. developed script for analyzing TCR sequence data; A.A.S., A.X., and E.C. performed *in vitro* TCR transduction experiments; E.S. performed *in vitro* cell culture; K.R.S. and M.K.-D. recruited T1D patients and collected blood samples from them. R.B. provided help on Rhapsody related experiments; K.-Y.M. and N.J. wrote the manuscript with help from all co-authors.

## Data Availability

All TCR and peptide information are in the Supplementary Tables. Raw sequencing data are being uploaded to dbGaP.

## Code Availability

Custom analysis code are being uploaded to Github.

