## Supplemental figures and tables for "High-Throughput and High-Dimensional Single Cell Analysis of Antigen-Specific CD8^+^ T cells"

[#]Current affiliation: Pfizer, China

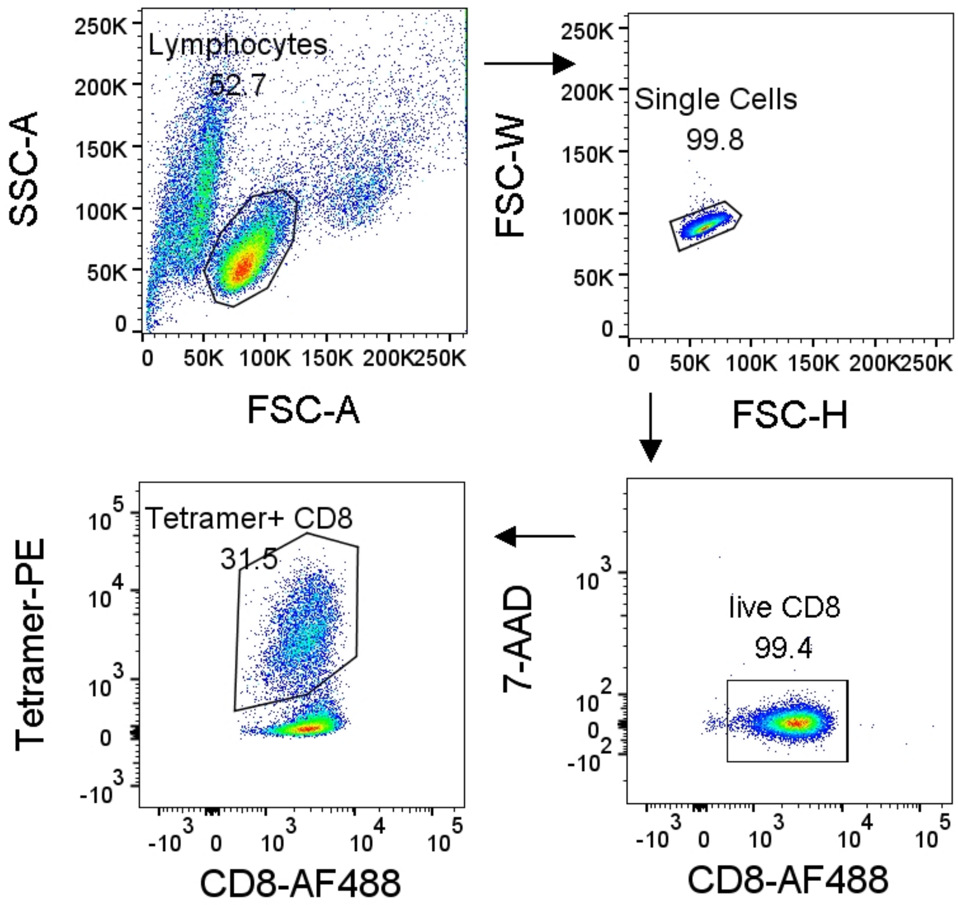


**Supplementary Fig. S1.** Gating strategy for sorting tetramer$^+$ CD8$^+$ T cells from a mixture of pMHC tetramer sorted polyclonal T cells cultured *in vitro*.

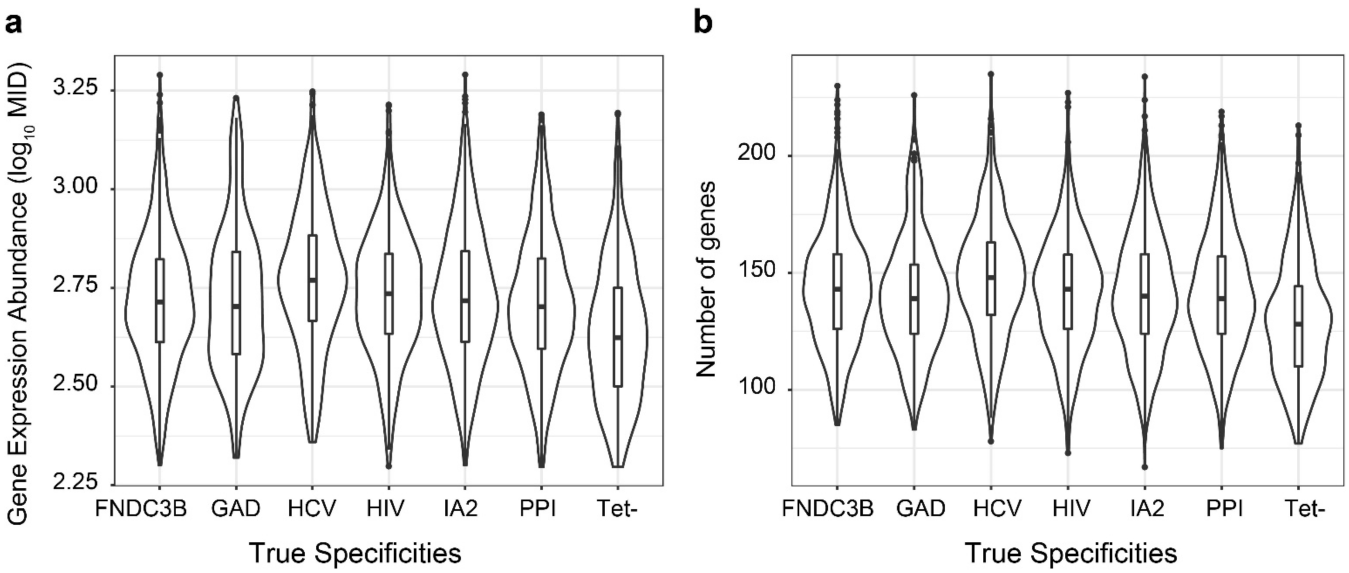


**Supplementary Fig. S2.** Distribution of (**a**) mRNA counts ($\log_{10}$) and (**b**) number of detected genes per cell among different antigen specific T cell populations.

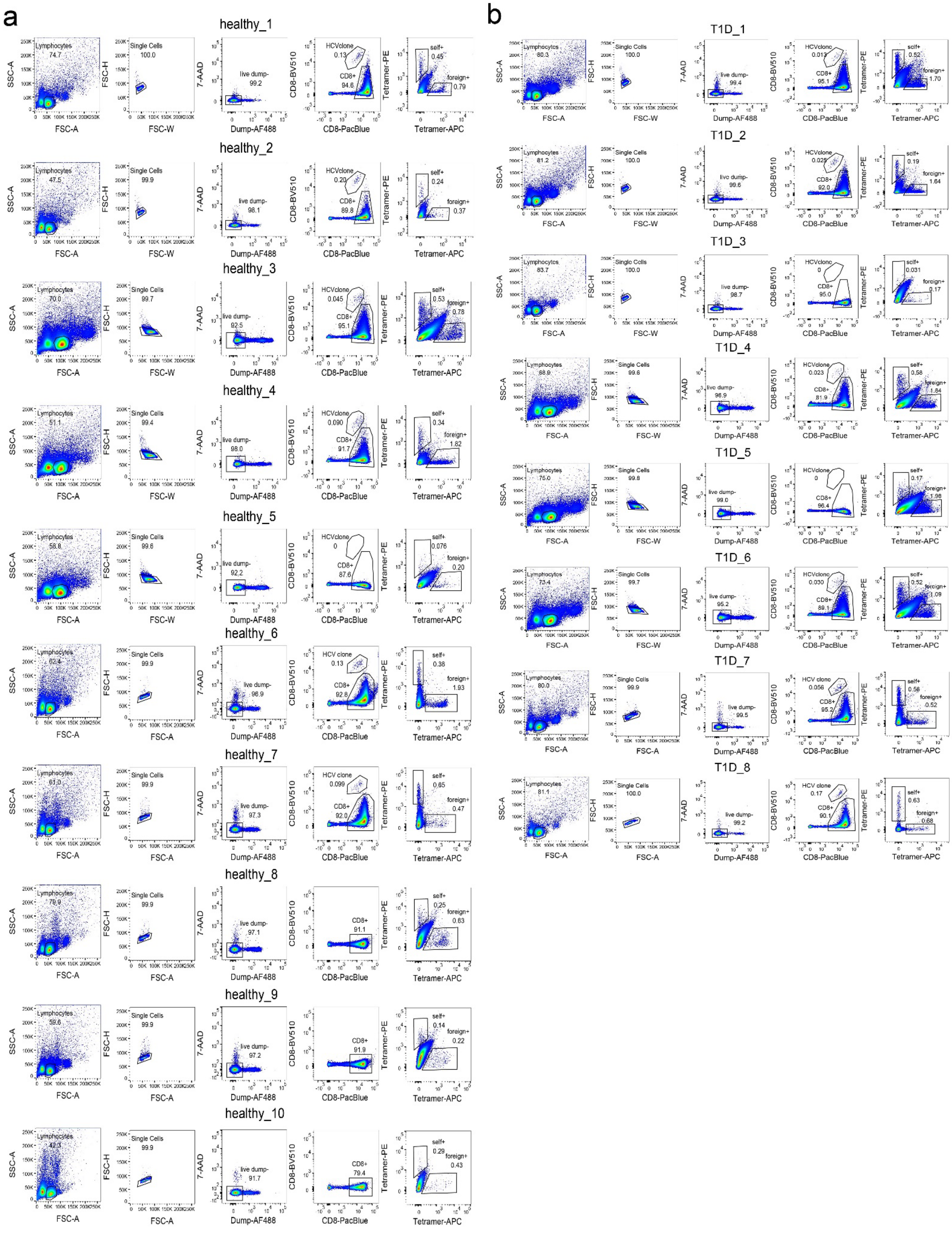

a
b
healthy_1
Lymphocytes 74.7
Single Cells 100.0
live dump- 99.2
HCVclone 0.13
CD8+ 94.6
self+ 0.45
foreign+ 0.79
healthy_2
Lymphocytes 47.5
Single Cells 99.9
live dump- 98.1
HCVclone 0.20
CD8+ 89.8
self+ 0.24
foreign+ 0.37
healthy_3
Lymphocytes 70.0
Single Cells 99.7
live dump- 92.5
HCVclone 0.045
CD8+ 95.1
self+ 0.53
foreign+ 0.78
healthy_4
Lymphocytes 51.1
Single Cells 99.4
live dump- 98.0
HCVclone 0.090
CD8+ 91.7
self+ 0.34
foreign+ 1.82
healthy_5
Lymphocytes 58.8
Single Cells 99.6
live dump- 92.2
HCVclone 0
CD8+ 87.6
self+ 0.076
foreign+ 0.20
healthy_6
Lymphocytes 62.4
Single Cells 99.9
live dump- 96.9
HCV clone 0.13
CD8+ 92.8
self+ 0.38
foreign+ 1.93
healthy_7
Lymphocytes 61.0
Single Cells 99.9
live dump- 97.3
HCV clone 0.099
CD8+ 92.0
self+ 0.65
foreign+ 0.47
healthy_8
Lymphocytes 79.9
Single Cells 99.9
live dump- 97.1
CD8+ 91.1
self+ 0.25
foreign+ 0.63
healthy_9
Lymphocytes 59.6
Single Cells 99.9
live dump- 97.2
CD8+ 91.9
self+ 0.14
foreign+ 0.22
healthy_10
Lymphocytes 42.3
Single Cells 99.9
live dump- 91.7
CD8+ 79.4
self+ 0.29
foreign+ 0.43
T1D_1
Lymphocytes 80.3
Single Cells 100.0
live dump- 99.4
HCVclone 0.013
CD8+ 95.1
self+ 0.52
foreign+ 1.70
T1D_2
Lymphocytes 81.2
Single Cells 100.0
live dump- 99.6
HCVclone 0.025
CD8+ 92.0
self+ 0.19
foreign+ 1.64
T1D_3
Lymphocytes 83.7
Single Cells 100.0
live dump- 98.7
HCVclone 0
CD8+ 95.0
self+ 0.031
foreign+ 0.17
T1D_4
Lymphocytes 88.9
Single Cells 99.6
live dump- 96.9
HCVclone 0.023
CD8+ 81.9
self+ 0.58
foreign+ 1.84
T1D_5
Lymphocytes 75.0
Single Cells 99.8
live dump- 99.0
HCVclone 0
CD8+ 96.4
self+ 0.17
foreign+ 1.98
T1D_6
Lymphocytes 73.4
Single Cells 99.7
live dump- 95.2
HCVclone 0.030
CD8+ 89.1
self+ 0.52
foreign+ 1.09
T1D_7
Lymphocytes 80.0
Single Cells 99.9
live dump- 99.5
HCV clone 0.056
CD8+ 95.2
self+ 0.56
foreign+ 0.52
T1D_8
Lymphocytes 81.1
Single Cells 100.0
live dump- 99.2
HCV clone 0.17
CD8+ 90.1
self+ 0.63
foreign+ 0.68
SSC-A
FSC-A
FSC-H
FSC-W
7-AAD
Dump-AF488
CD8-BV510
CD8-PacBlue
Tetramer-PE
Tetramer-APC

**Supplementary Fig. S3.** Gating strategy to sort tetramer$^+$ T cells from primary CD8$^+$ T cells enriched from PBMCs of (**a**) healthy and (**b**) T1D donors. Cultured HCV antigen-specific clone was spiked in primary CD8$^+$ T cells whose HLA alleles include A02:01 and a different fluorescently labeled CD8$^+$ antibody was used to pre-stain HCV antigen-specific clone.

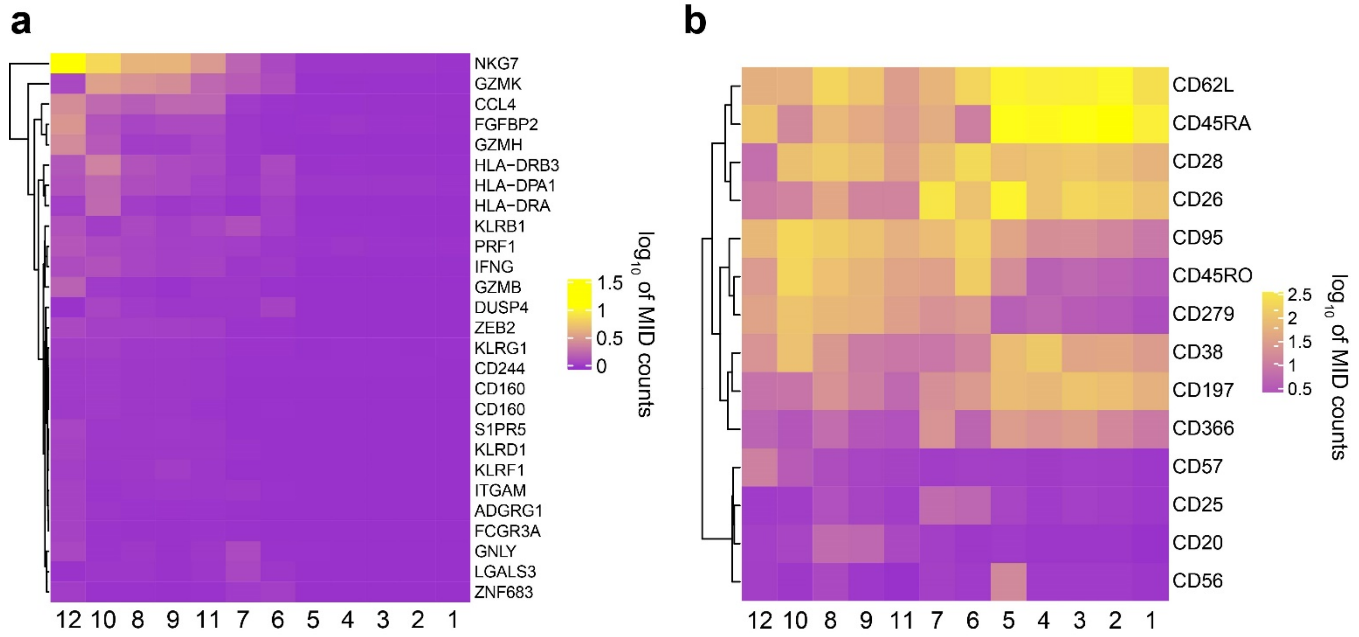


**Supplementary Fig. S4.** Absolute expression ($log_{10}$ of MID counts) of differentially expressed (**a**) genes and (**b**) surface-proteins among different clusters. Genes and surface-protein plotted here are the same set as in **Figure 2c**.

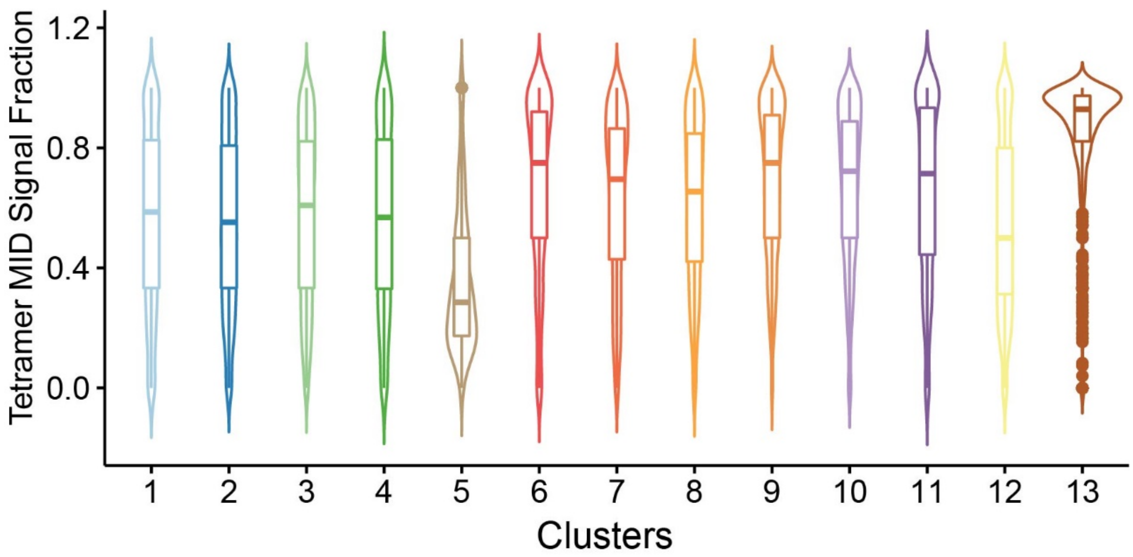


**Supplementary Fig. S5.** The distribution of tetramer DNA barcode MID signal fraction among different phenotype clusters combined from all donors. For each cell, tetramer MID signal fraction was defined as the fraction of cumulative MID count from putative binding antigens over cumulative MID count from all bound antigens (**See methods**).

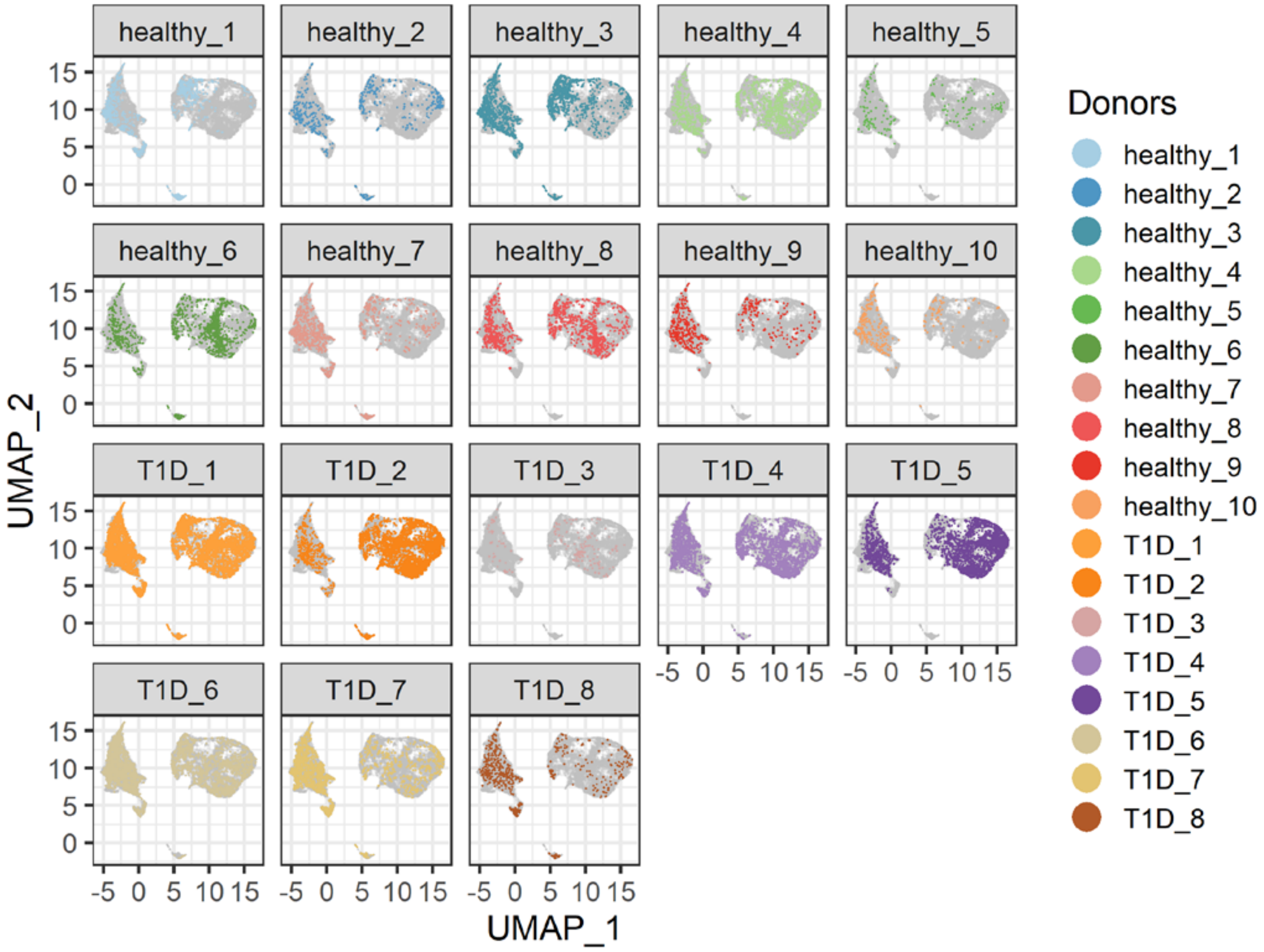


**Supplementary Fig. S6.** UMAP projection of single cells among different donors. Grey dots represent all cells and colored dots are cells from different chips.

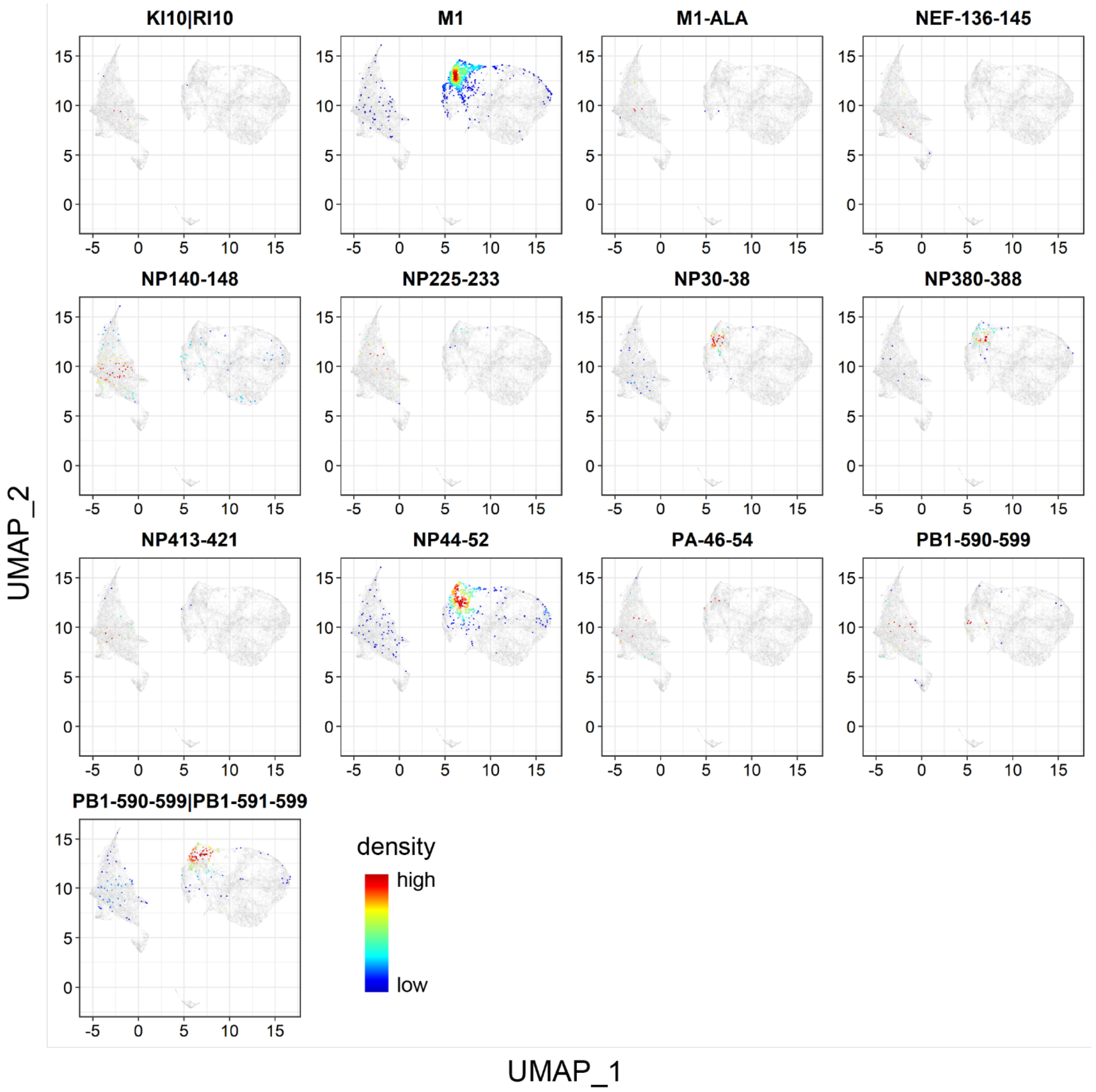


**Supplementary Fig. S7.** Density heatmap of the projection of influenza specificic T cells on UMAP plot. Colors represent the local density of cells on the two-dimension space.

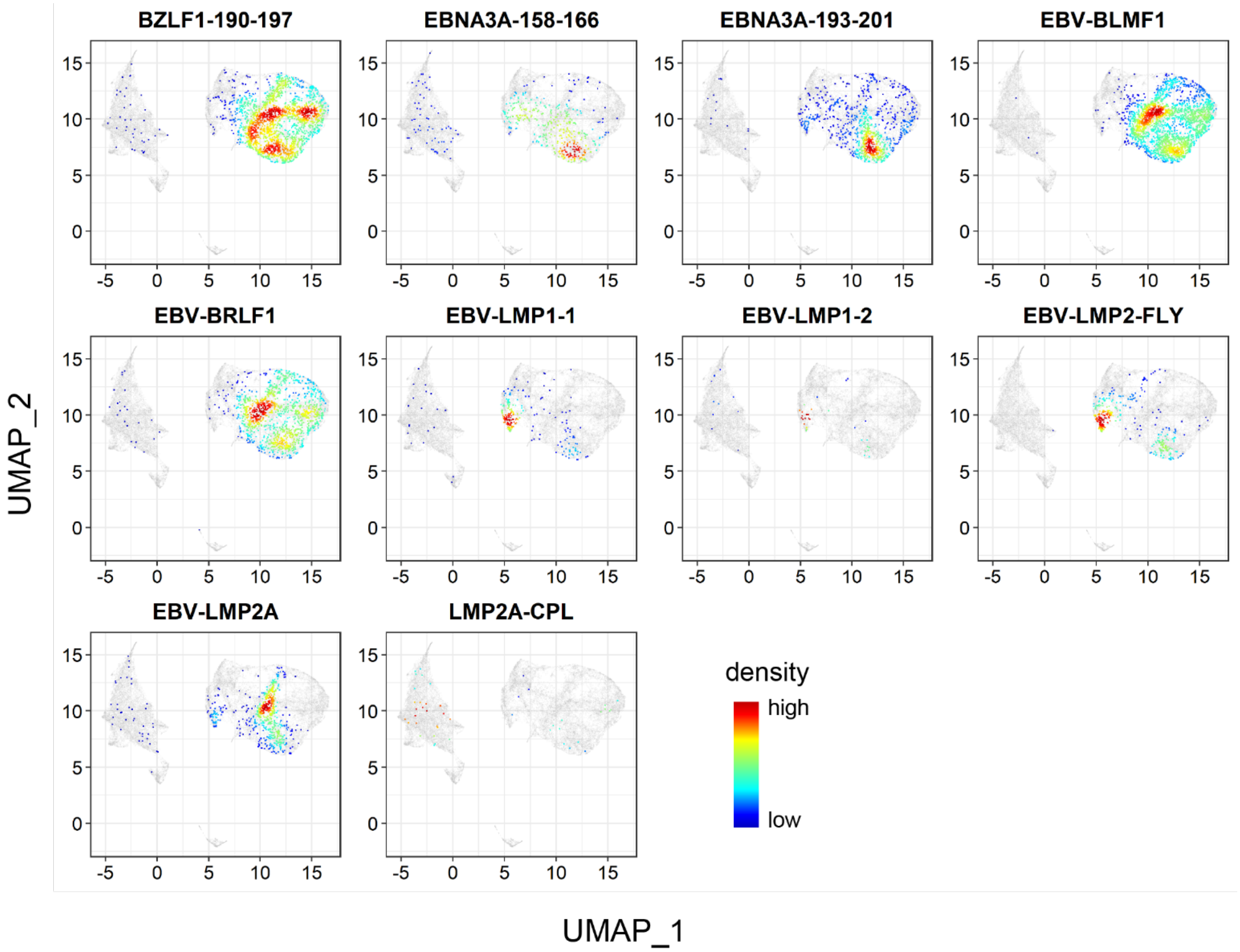


**Supplementary Fig. S8.** Density heatmap of the projection of EBV specificic T cells on UMAP plot. Colors represent the local density of cells on the two-dimension space.

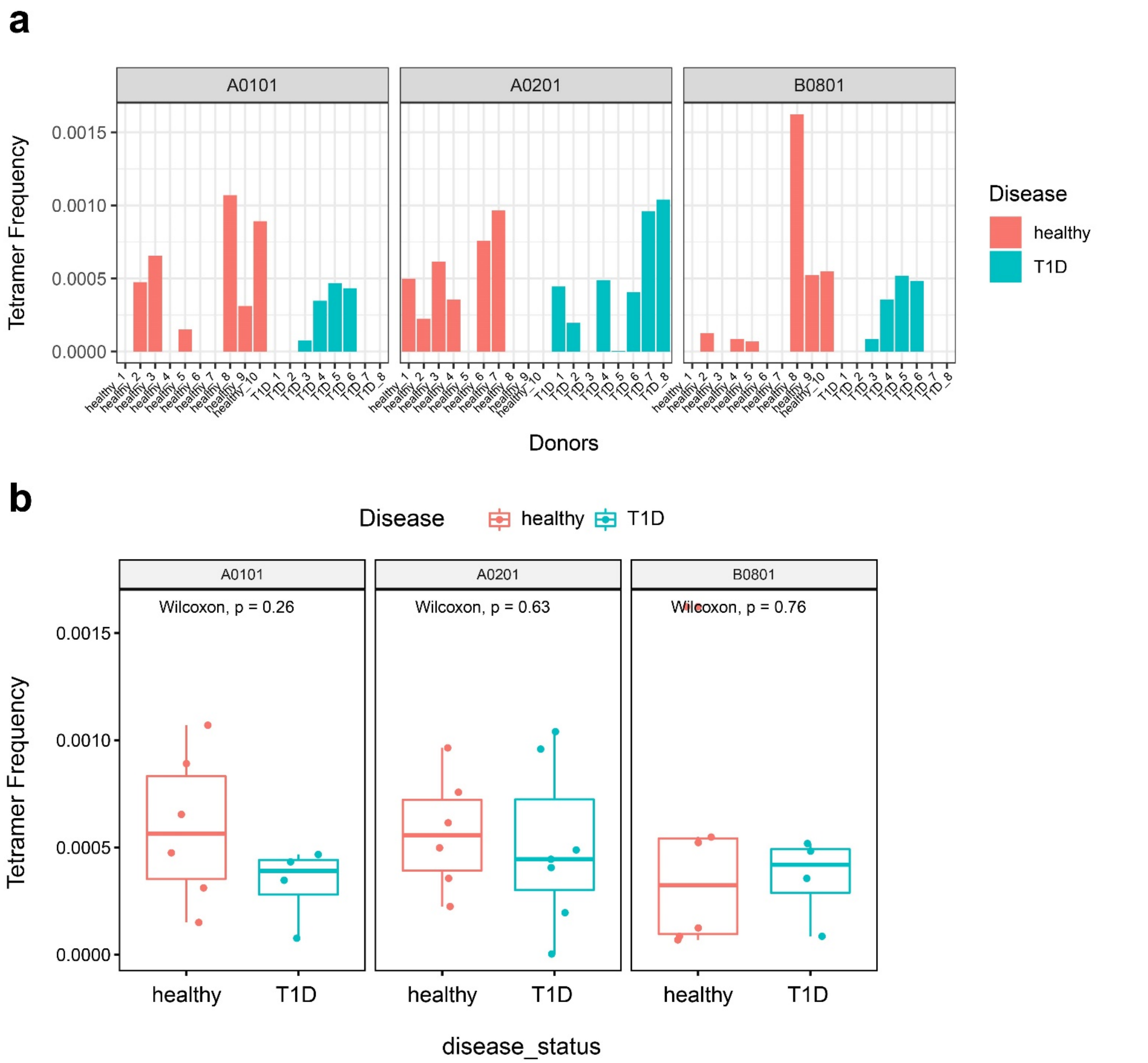


**Supplementary Fig. S9.** Frequency of total T1D autoantigen-specific CD8$^+$ T cells in healthy subjects and T1D patients. **(a**). Frequency of T1D autoantigen tetramer$^+$ CD8$^+$ T cells in different donors for various HLA alleles. (**b**). Comparison of total T1D autoantigen tetramer$^+$ CD8$^+$ T cells between healthy and T1D donors for various HLA alleles. Wilcoxon non-parametric test was performed.

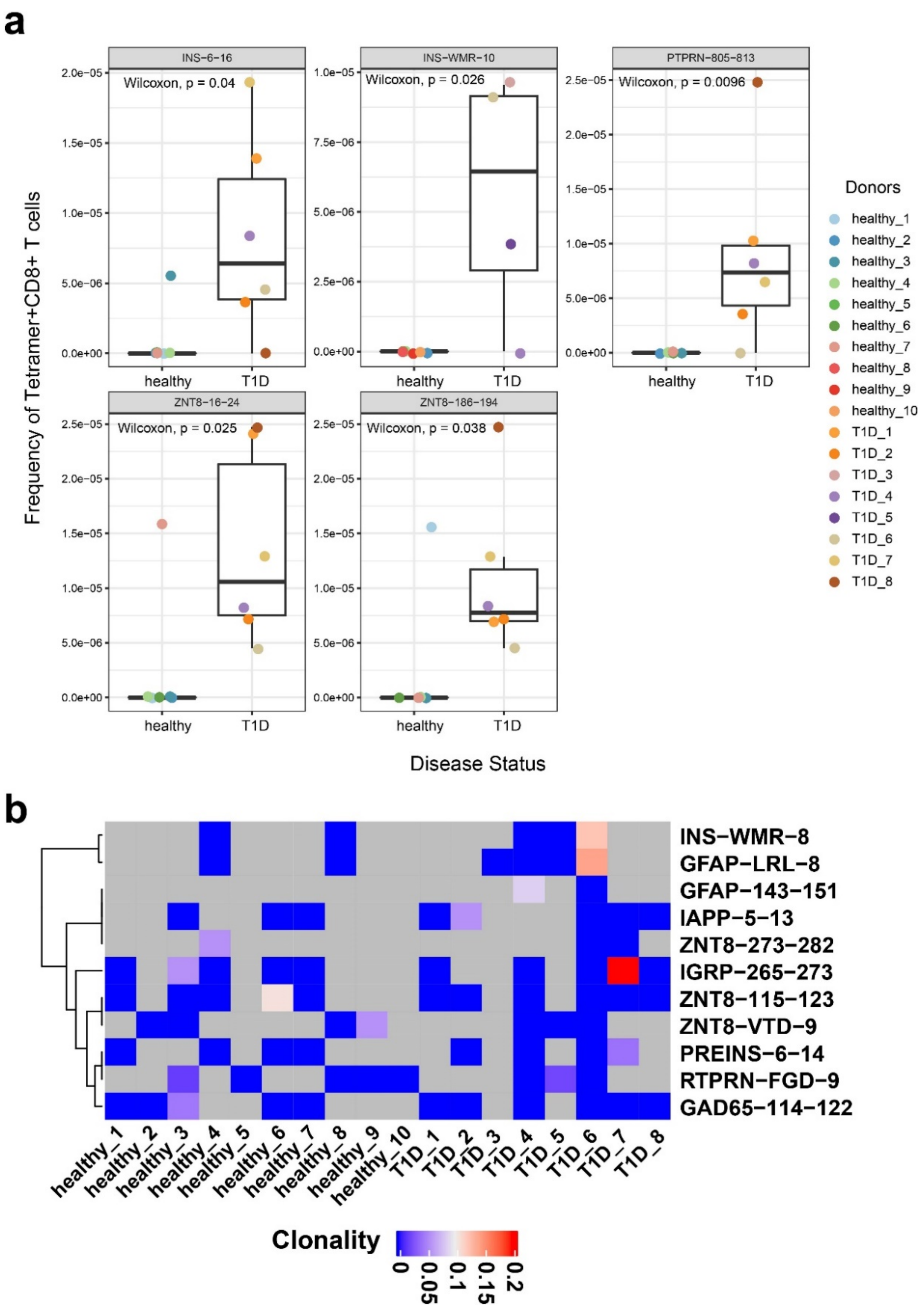


**Supplementary Fig. S10.** T1D autoantigens with different antigen-specific CD8$^{+}$ T cell frequencies and clonality between healthy subjects and T1D patients. **(a)**. Five T1D autoantigens were identified to have significant higher frequency of antigen specific T cells in peripheral blood when MID negative threshold was set to 15. Wilcoxon non-parametric test was performed. (**b**). TCR clonality heatmap of T1D antigenic specific T cells for each antigen/donor combination. Grey, no T cells were detected.

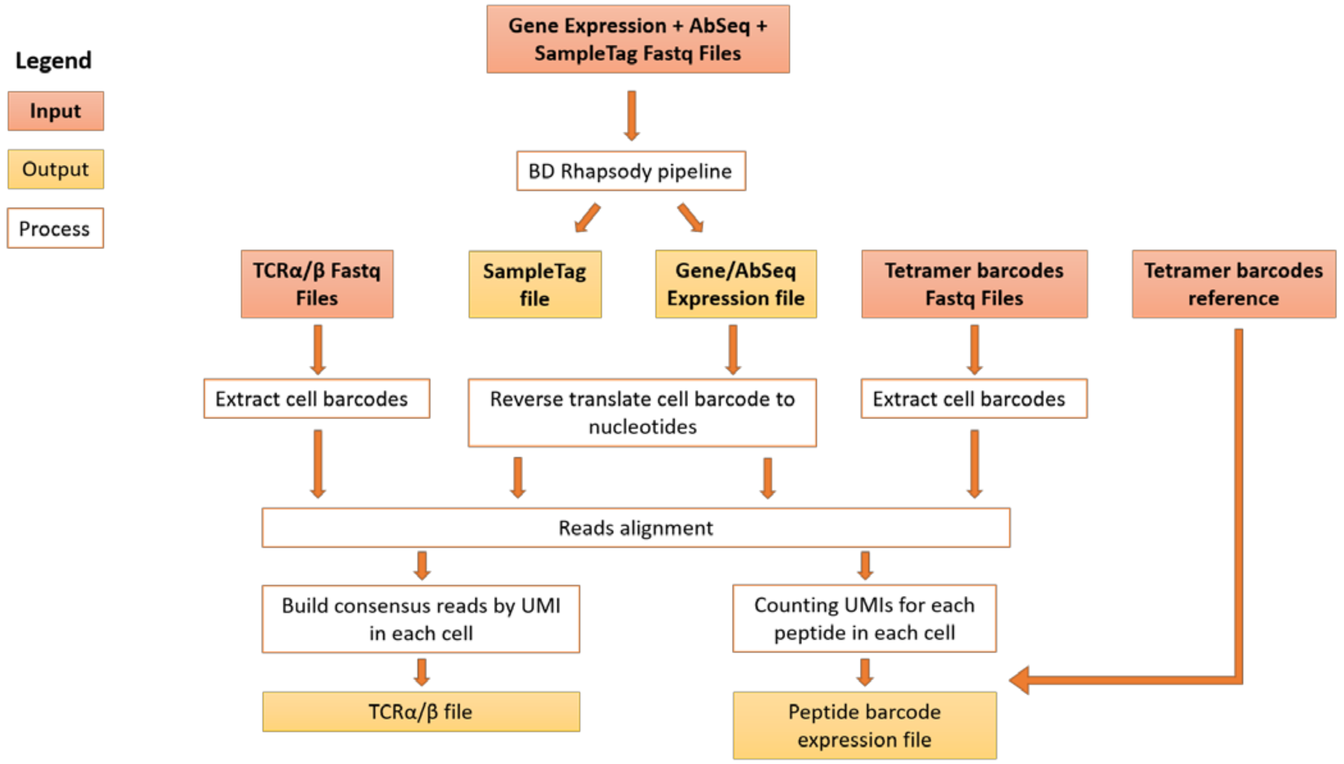


**Supplementary Fig. S11.** Diagrams illustrating the bioinformatic workflow to analyze target gene expression, AbSeq, SampleTag, tetramer DNA barcode, and TCR simultaneously for single cells.

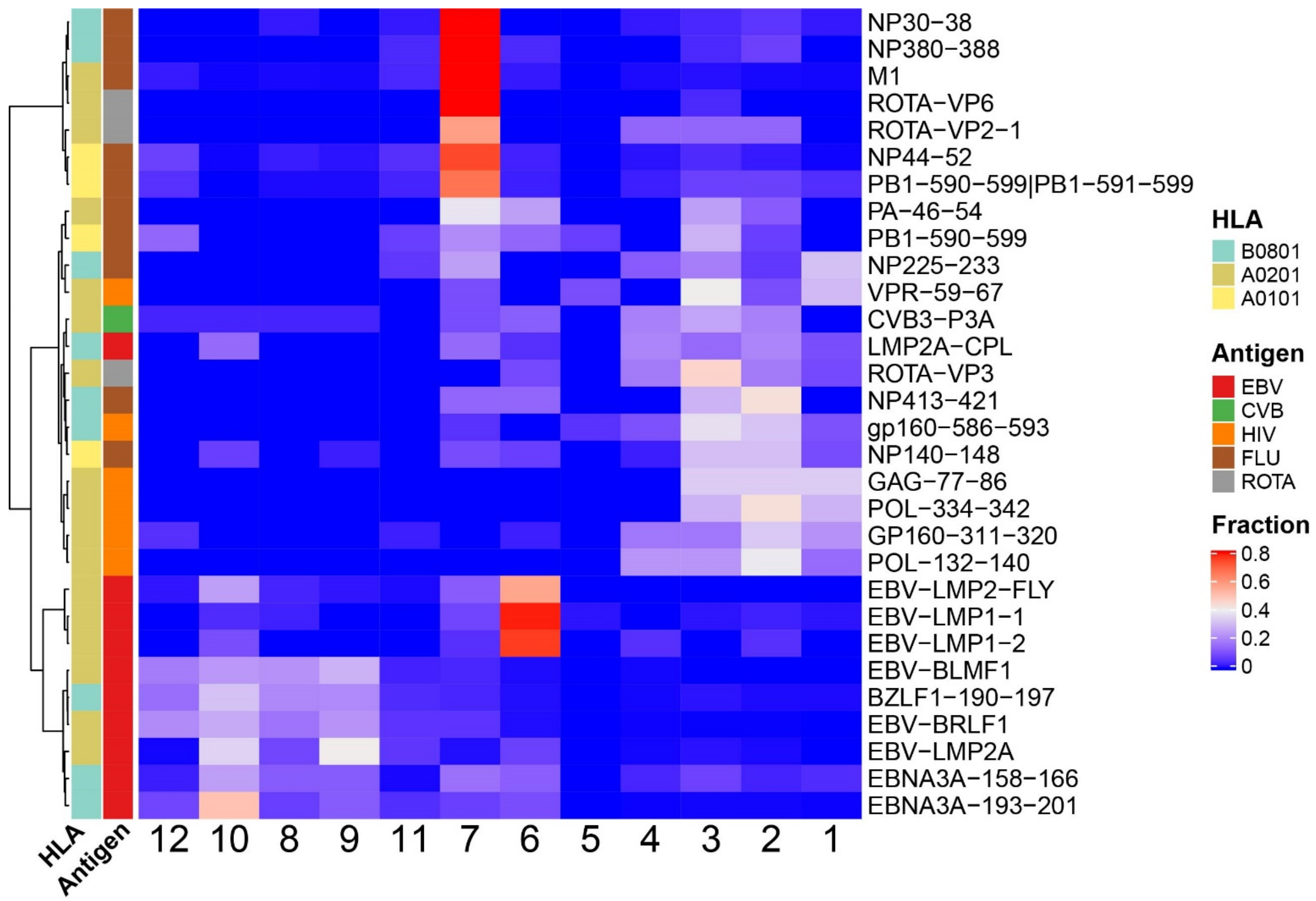


**Supplementary Fig. S12.** Distribution of viral antigen specific CD8⁺ T cells among 12 primary CD8⁺ T cells clusters in all 18 donors when tetramer MID negative threshold was set to 15 (**See methods**).

**Supplementary Table 1: Antigens used to sort and stimulate polyclonal CD8+ T cells.**

| Antigen | DNA barcode | Antigen Peptide Sequence | Gene | Category |
|---|---|---|---|---|
| IA2 | TTCATTGACTCTTACATCTGCCAGGTT | SLSPLQAEL | IA2 | self |
| GAD | TGGCTGTCTCTGCTGGTTCCGTTTGTT | VMNILLQYVV | GAD | self |
| PPI | GCTCTGTGGATGCGTCTGCTGCCGCTG | ALWMRLLPL | PPI | self |
| HCV | AAACTGGTTGCTCTGGGTATCAACGCTGTT | KLVALGINAV | HCV | foreign |
| HIV | TCTCTGTATAACACCGTGGCTACCCTG | LLNATDIAV | HIV | foreign |
| FNDC3B_WT | CTGCTGTGGAACGGTCCGATGGCTGTT | VVLSWAPPV | FNDC3B | self |
| FNDC3B_MUT | TGCCTGGGTGGTCTGCTGACGATGGTT | VVMSWAPPV | FNDC3B | foreign |
| Empty* | AACCTGGTTCCGATGGTTGCTACCGTT | Empty | Empty | Empty |

*Empty: Negative control without antigen loaded on pMHC.

**Supplementary Table 2: Sequencing metrics for all TetTCR-SeqHD experiments**

| | Clone Experiment | Chip1 | Chip2 |
|---|---|---|---|
| Number of Cells | 4533 | 11573 | 15007 |
| Total Gene Expression Reads (reads per cell) | 49333947 (10883) | 52038256 (4497) | 64831148 (4320) |
| Total AbSeq Reads (reads per cell) | N/A | 287599407 (24851) | 289247804 (19274) |
| Total Sampletag Reads (reads per cell) | N/A | 42824048 (3700) | 81079634 (5403) |
| Total Tetramer Reads (reads per cell) | 78190817 (17249) | 126331571 (10916) | 148948212 (9925) |
| Total TCRa Reads (reads per cell) | 4176402 (921) | 43854843 (3789) | 55898962 (3725) |
| Total TCRb Reads (reads per cell) | 2032361 (448) | 39622085 (3424) | 42322117 (2820) |
| Total Reads (reads per cell) | 133733527 (29502) | 592270210 (51177) | 682327877 (45467) |

Note: Number in the bracket indicates the average number of reads per cell

| Chip3 | Chip4 |
|---|---|
| 3983 | 4605 |
| 27174272 (6823) | 20377883 (4425) |
| 106951767 (26852) | 95129527 (20658) |
| 19491789 (4894) | 20631061 (4480) |
| 60396499 (15164) | 37430622 (8128) |
| 16996596 (4267) | 11570756 (2513) |
| 13223672 (3320) | 10682431 (2320) |
| 244234595 (61319) | 195822280 (42524) |

**Supplementary Table 3: Reference sequences of TCRβ for polyclonal CD8+ T cells**

| Clone | Clone Count | AA sequence of CDR3 | Gene |
|---|---|---|---|
| IA2_45055 | 45055 | CSAIVLAGRTDTQYF | IA2 |
| IA2_39321 | 39321 | CAISQGDTEAFF | IA2 |
| IA2_3102 | 3102 | CSVPGTEGNEQFF | IA2 |
| IA2_68 | 68 | CASTVGGTHDTDTQYF | IA2 |
| IA2_24 | 24 | CSVGTGGNYGYTF | IA2 |
| IA2_13 | 13 | CASSFSTCSANYGYTF | IA2 |
| PPI_6717 | 6717 | CASSSGAEAFF | PPI |
| PPI_4136 | 4136 | CASSDVADSRPGDEQYF | PPI |
| PPI_3269 | 3269 | CASSSPLAGRPNEQFF | PPI |
| PPI_270 | 270 | CASSYLAGGADSSTQYF | PPI |
| PPI_127 | 127 | CASKGETGGWTGELFF | PPI |
| PPI_21 | 21 | CASSQTFEPNGFNEQFF | PPI |
| PPI_5 | 5 | CASSLRGTTYEQYF | PPI |
| GAD_68701 | 68701 | CASSQDVAGNYGYTF | GAD |
| GAD_6485 | 6485 | CASTKGDSYGYTF | GAD |
| GAD_1706 | 1706 | CASSYLSLQGTSYNEQFF | GAD |
| GAD_1260 | 1260 | CASSDVGGSEQFF | GAD |
| GAD_1062 | 1062 | CASRFLGTEAFF | GAD |
| GAD_862 | 862 | CASSDLLGSPNTEAFF | GAD |
| GAD_434 | 434 | CASWTDEAFF | GAD |
| GAD_418 | 418 | CASSEGGLAGLTDTQYF | GAD |
| GAD_66 | 66 | CASSELGSPVEKLFF | GAD |
| GAD_33 | 33 | CASSLAGDVWDNTEAFF | GAD |
| GAD_15 | 15 | CASSARTSGRATYSYNEQFF | GAD |
| HCV_58236 | 58236 | CASSAMGSGNTIYF | HCV |
| HCV_38531 | 38531 | CSARDRDEKLFF | HCV |
| HCV_703 | 703 | CASTHFEDEQFF | HCV |
| HCV_165 | 165 | CASSIARDRVSNQPQHF | HCV |
| HCV_86 | 86 | CSASGQGADQPQHF | HCV |
| HCV_15 | 15 | CASATSSYEQYF | HCV |
| HCV_4 | 4 | CASSSTGTGGNEQFF | HCV |
| HIV_33798 | 33798 | CASSGRQGGYTF | HIV |
| HIV_21405 | 21405 | CSASAANTGELFF | HIV |
| HIV_6571 | 6571 | CASSLSRFGSYEQYF | HIV |
| HIV_122 | 122 | CASMKAGQETQYF | HIV |
| HIV_118 | 118 | CASSFGTGGYEQYF | HIV |
| FNDC3B-CROSS_62231 | 62231 | CSALEVGPQEYF | FNDC3B |
| FNDC3B-CROSS_54891 | 54891 | CASSFGPTGYEQYF | FNDC3B |
| FNDC3B-CROSS_2208 | 2208 | CSPRGPYNEQFF | FNDC3B |
| FNDC3B-CROSS_1010 | 1010 | CASSLNGPSYNEQFF | FNDC3B |
| FNDC3B-CROSS_739 | 739 | CASSLGGGSPLHF | FNDC3B |
| FNDC3B-CROSS_617 | 617 | CASSIGPSYEQYF | FNDC3B |
| FNDC3B-CROSS_140 | 140 | CASTDNPAGGGYGYTF | FNDC3B |
| FNDC3B-CROSS_81 | 81 | CASRYLRGSSYEQYF | FNDC3B |
| FNDC3B-CROSS_68 | 68 | CASSKGLPTYNEQFF | FNDC3B |

| | | | |
|---|---|---|---|
| FNDC3B-CROSS_13 | 13 | CASSFATGVGSYGYTF | FNDC3B |

**Supplementary Table 4: Endogenouse and foreign antigens used in TetTCR-SeqHD experin**

| | Antigen | DNA barcode |
|---|---|---|
| 1 | ADH-79-88 | CGTCAGTTCGGTCCGGACTGGATCGTTGCT |
| 2 | DUF5119-124-133 | ATGGTTTGGGGTCCGGACCCGCTGTACGTT |
| 3 | GFAP-143-151 | AACCTGGCTCAGGACCTGGCTACCGTT |
| 4 | GFAP-214-222 | CAGCTGGCTCGTCAGCAGGTTCACGTT |
| 5 | GAD65-110-118 | TTCCTGCAGGACGTTATGAACATCCTG |
| 6 | GAD65-141-149 | CTGCTGCAGGAATACAACTGGGAACTG |
| 7 | GAD65-536-545 | CGTATGATGGAATACGGTACCACCATGGTT |
| 8 | GAD65-114-123 | GTTATGAACATCCTGCTGCAGTACGTTGTT |
| 9 | GAD65-114-122 | GTTATGAACATCCTGCTGCAGTACGTT |
| 10 | GAD65-476-484 | GAACTGGCTGAATACCTGTACAACATC |
| 11 | GAD65-159-167 | ATCCTGATGCACTGCCAGACCACCCTG |
| 12 | INSDRIP-1-9 | ATGCTGTACCAGCACCTGCTGCCGCTG |
| 13 | INS-78-87 | GGTATCGTTGAACAGTGCTGCACCTCTATC |
| 14 | INS-2-11 | GCTCTGTGGATGCGTCTGCTGCCGCTGCTG |
| 15 | INS-14-23 | CTGGCTCTGTGGGGTCCGGACCCGGCTGCT |
| 16 | INS-6-16 | CGTCTGCTGCCGCTGCTGGCTCTGCTGGCTCTG |
| 17 | PREINS-2-10 | GCTCTGTGGATGCGTCTGCTGCCGCTG |
| 18 | PREINS-34-42 | CACCTGGTTGAAGCTCTGTACCTGGTT |
| 19 | PREINS-85-94 | TCTCTGCAGAAACGTGGTATCGTTGAACAG |
| 20 | PREINS-76-84 | TCTCTGCAGCCGCTGGCTCTGGAAGGT |
| 21 | PREINS-101-109 | TCTCTGTACCAGCTGGAAAACTACTGC |
| 22 | PREINS-42-51 | GTTTGCGGTGAACGTGGTTTCTTCTACACC |
| 23 | PREINS-15-24 | GCTCTGTGGGGTCCGGACCCGGCTGCTGCT |
| 24 | PREINS-6-14 | CGTCTGCTGCCGCTGCTGGCTCTGCTG |
| 25 | PREINS-17-24 | TGGGGTCCGGACCCGGCTGCTGCT |
| 26 | IAPP-9-17 | TTCCTGATCGTTCTGTCTGTTGCTCTG |
| 27 | IAPP-5-13 | AAACTGCAGGTTTTCCTGATCGTTCTG |
| 28 | IGRP-152-160-MLI | TTCCTGTGGTCTGTTTTCATGCTGATC |
| 29 | IGRP-215-223 | TTCCTGTTCGCTGTTGGTTTCTACCTG |
| 30 | IGRP-228-236 | CTGAACATCGACCTGCTGTGGTCTGTT |
| 31 | IGRP-265-273 | GTTCTGTTCGGTCTGGGTTTCGCTATC |
| 32 | IGRP-152-160-WLI | TTCCTGTGGTCTGTTTTCTGGCTGATC |
| 33 | IGRP-211-219 | AACCTGTTCCTGTTCCTGTTCGCTGTT |
| 34 | IGRP-222-230 | TACCTGCTGCTGCGTGTTCTGAACATC |
| 35 | PPI-29-38 | CACCTGTGCGGTTCTCACCTGGTTGAAGCT |
| 36 | PPI-33-42 | TCTCACCTGGTTGAAGCTCTGTACCTGGTT |
| 37 | PPI-30-39 | CTGTGCGGTTCTCACCTGGTTGAAGCTCTG |
| 38 | PTPRN-962-970 | GCTCTGACCGCTGTTGCTGAAGAAGTT |
| 39 | PTPRN-830-839 | TCTCTGTACCACGTTTACGAAGTTAACCTG |
| 40 | PTPRN-790-798 | ACCATCGCTGACTTCTGGCAGATGGTT |
| 41 | PTPRN-805-813 | GTTATCGTTATGCTGACCCCGCTGGTT |
| 42 | PTPRN-180-188 | CTGCTGCCGCCGCTGCTGGAACACCTG |
| 43 | PTPRN-482-490 | TCTCTGGCTGCTGGTGTTAAACTGCTG |
| 44 | PTPRN-172-180 | TCTCTGTCTCCGCTGCAGGCTGAACTG |

| | | |
|---|---|---|
| 45 | PTPRN-797-805 | ATGGTTTGGGAATCTGGTTGCACCGTT |
| 46 | ZNT8 | GTTATGATCATCGTTTCTTCTCTGGCTGTT |
| 47 | ZNT8-221-229 | GCTCTGGGTGACCTGTTCCAGTCTATC |
| 48 | ZNT8-110-118 | GACCTGACCTCTTTCCTGCTGTCTCTG |
| 49 | ZNT8-140-148 | GAAATCCTGGGTGCTCTGCTGTCTATC |
| 50 | ZNT8-114-123 | TTCCTGCTGTCTCTGTTCTCTCTGTGGCTG |
| 51 | ZNT8-291-300 | ATCCTGGCTGTTGACGGTGTTCTGTCTGTT |
| 52 | ZNT8-141-149 | ATCCTGGGTGCTCTGCTGTCTATCCTG |
| 53 | ZNT8-266-274 | ATCCTGAAAGACTTCTCTATCCTGCTG |
| 54 | ZNT8-314-322 | ATCCTGTCTGCTCACGTTGCTACCGCT |
| 55 | ZNT8-107-115 | CTGCTGATCGACCTGACCTCTTTCCTG |
| 56 | ZNT8-273-282 | CTGCTGATGGAAGGTGTTCCGAAATCTCTG |
| 57 | ZNT8-228-236 | TCTATCTCTGTTCTGATCTCTGCTCTG |
| 58 | ZNT8-281-290 | TCTCTGAACTACTCTGGTGTTAAAGAACTG |
| 59 | ZNT8-299-308 | TCTGTTCACTCTCTGCACATCTGGTCTCTG |
| 60 | ZNT8-153-161 | GTTGTTACCGGTGTTCTGGTTTACCTG |
| 61 | ZNT8-253-261 | TTCATCTTCTCTATCCTGGTTCTGGCT |
| 62 | ZNT8-173-181 | ATCCAGGCTACCGTTATGATCATCGTT |
| 63 | ZNT8-16-24 | AAAATGTACGCTTTCACCCTGGAATCT |
| 64 | ZNT8-280-288 | AAATCTCTGAACTACTCTGGTGTTAAA |
| 65 | ZNT8-293-300 | CTGGCTGTTGACGGTGTTCTGTCTGTT |
| 66 | ZNT8-115-123 | CTGCTGTCTCTGTTCTCTCTGTGGCTG |
| 67 | ZNT8-165-173 | CGTCTGCTGTACCCGGACTACCAGATC |
| 68 | ZNT8-343-351 | ACCATGCACTCTCTGACCATCCAGATG |
| 69 | ZNT8-186-194 | GTTGCTGCTAACATCGTTCTGACCGTT |
| 70 | ZNT8-200-208 | TGCCTGGGTCACAACCACAAAGAAGTT |
| 71 | ZNT8-245-254 | AAAATCGCTGACCCGATCTGCACCTTCATC |
| 72 | ZNT8-16-25 | AAAATGTACGCTTTCACCCTGGAATCTGTT |
| 73 | ZNT8-107-116 | CTGCTGATCGACCTGACCTCTTTCCTGCTG |
| 74 | ZNT8-145-153 | CTGCTGTCTATCCTGTGCATCTGGGTT |
| 75 | GAG-77-86 | TCTCTGTACAACACCGTTGCTACCCTGTAC |
| 76 | GAG-433-442 | TTCCTGGGTAAAATCTGGCCGTCTTACAAA |
| 77 | POL-132-140 | CTGGTTGGTCCGACCCCGGTTAACATC |
| 78 | POL-188-196 | GCTCTGGTTGAAATCTGCACCGAAATG |
| 79 | POL-334-342 | GTTATCTACCAGTACATGGACGACCTG |
| 80 | POL-464-472 | ATCCTGAAAGAACCGGTTCACGGTGTT |
| 81 | VPR-59-67 | GCTATCATCCGTATCCTGCAGCAGCTG |
| 82 | GP160-311-320 | CGTGGTCCGGGTCGTGCTTTCGTTACCATC |
| 83 | GP160-813-822 | TCTCTGCTGAACGCTACCGACATCGCTGTT |
| 84 | GP160-814-822 | CTGCTGAACGCTACCGACATCGCTGTT |
| 85 | NEF-136-145 | CCGCTGACCTTCGGTTGGTGCTACAAACTG |
| 86 | NEF-180-189 | GTTCTGGAATGGCGTTTCGACTCTCGTCTG |
| 87 | M1 | GGTATCCTGGGTTTCGTTTTCACCCTG |
| 88 | KI10 | AAACTGTACCAGAACCCGACCACCTACATC |
| 89 | RI10 | CGTCTGTACCAGAACCCGACCACCTACATC |
| 90 | NS1-122-130 | GCTATCATGGACAAAAACATCATCCTG |
| 91 | PA-46-54 | TTCATGTACTCTGACTTCCACTTCATC |

| | | |
|---|---|---|
| 92 | HCVNS3-1406-1415 | AAACTGGTTGCTCTGGGTATCAACGCTGTT |
| 93 | HCVNS4B-1807-1816 | CTGCTGTTCAACATCCTGGGTGGTTGGGTT |
| 94 | HCVNS3-1073-1081 | TGCATCAACGGTGTTTGCTGGACCGTT |
| 95 | HCV-35-44 | TACCTGCTGCCGCGTCGTGGTCCGCGTCTG |
| 96 | EBV-LMP2A | TGCCTGGGTGGTCTGCTGACCATGGTT |
| 97 | EBV-LMP1-2 | TACCTGCAGCAGAACTGGTGGACCCTG |
| 98 | EBV-LMP1-1 | TACCTGCTGGAAATGCTGTGGCGTCTG |
| 99 | EBV-BRLF1 | TACGTTCTGGACCACCTGATCGTTGTT |
| 100 | EBV-BLMF1 | GGTCTGTGCACCCTGGTTGCTATGCTG |
| 101 | EBV-LMP2-FLY | TTCCTGTACGCTCTGGCTCTGCTGCTG |
| 102 | ROTA-VP3 | TACCTGCTGCCGGGTTGGAAACTG |
| 103 | ROTA-VP2-1 | TCTCTGATCTCTGGTATGTGGCTGCTG |
| 104 | ROTA-VP6 | ACCCTGCTGGCTAACGTTACCGCTGTT |
| 105 | CVB4-P2C | GAAGTTAAAGAAAAACACGAATTCCTG |
| 106 | CVB3-P3C | ATCCTGATGAACGACCAGGAAGTTGGTGTT |
| 107 | CVB3-P3A | GGTATCATCTACATCATCTACAAACTG |
| 108 | MART1-26-35 | GAAGCTGCTGGTATCGGTATCCTGACCGTT |
| 109 | MART1-A27L | GAACTGGCTGGTATCGGTATCCTGACCGTT |
| 110 | MART1-ALA | GCTCTGGCTGGTATCGGTATCCTGACCGTT |
| 111 | MART1-27-35 | GCTGCTGGTATCGGTATCCTGACCGTT |
| 112 | MART1-A28L | GCTCTGGGTATCGGTATCCTGACCGTT |
| 113 | PGT-178 | CTGCTGGCTGGTATCGGTACCGTTCCGATC |
| 114 | HCV-K1Y | TACCTGGTTGCTCTGGGTATCAACGCTGTT |
| 115 | HCV-K1YI7V | TACCTGGTTGCTCTGGGTGTTAACGCTGTT |
| 116 | HCV-A9N | AAACTGGTTGCTCTGGGTATCAACAACGTT |
| 117 | HCV-K1S | TCTCTGGTTGCTCTGGGTATCAACGCTGTT |
| 118 | HCV-L2I | AAAATCGTTGCTCTGGGTATCAACGCTGTT |
| 119 | INS-CTS-8 | TGCACCTCTATCTGCTCTCTGTAC |
| 120 | INS-CGS-10 | TGCGGTTCTCACCTGGTTGAAGCTCTGTAC |
| 121 | INS-GSH-9 | GGTTCTCACCTGGTTGAAGCTCTGTAC |
| 122 | GAD2-CLE-9 | TGCCTGGAACTGGCTGAATACCTGTAC |
| 123 | GAD2-STA-10 | TCTACCGCTAACACCAACATGTTCACCTAC |
| 124 | GAD2-KCL-10 | AAATGCCTGGAACTGGCTGAATACCTGTAC |
| 125 | GAD2-QQD-10 | CAGCAGGACAAACACTACGACCTGTCTTAC |
| 126 | GAD2-VSA-10 | GTTTCTGCTACCGCTGGTACCACCGTTTAC |
| 127 | GAD2-STK-9 | TCTACCAAAGTTATCGACTTCCACTAC |
| 128 | ZNT8-YLA-9 | TACCTGGCTTGCGAACGTCTGCTGTAC |
| 129 | ZNT8-VTD-9 | GTTACCGACGCTGCTCACCTGCTGATC |
| 130 | ZNT8-PTE-9 | CCGACCGAAAAAGGTGCTAACGAATAC |
| 131 | ZNT8-LID-9 | CTGATCGACCTGACCTCTTTCCTGCTG |
| 132 | ZNT8-KPT-10 | AAACCGACCGAAAAAGGTGCTAACGAATAC |
| 133 | ZNT8-VVT-10 | GTTGTTACCGACGCTGCTCACCTGCTGATC |
| 134 | ZNT8-LTS-9 | CTGACCTCTTTCCTGCTGTCTCTGTTC |
| 135 | IAPP-STN-9 | TCTACCAACGTTGGTTCTAACACCTAC |
| 136 | IAPP-SST-10 | TCTTCTACCAACGTTGGTTCTAACACCTAC |
| 137 | G6PC2-LTS-10 | CTGACCTCTCTGACCATCCTGCAGCTGTAC |
| 138 | G6PC2-PTH-9 | CCGACCCACGAAGAACACCTGTTCTAC |

| | | |
|---|---|---|
| 139 | G6PC2-IPT-10 | ATCCCGACCCACGAAGAACACCTGTTCTAC |
| 140 | G6PC2-TSL-9 | ACCTCTCTGACCATCCTGCAGCTGTAC |
| 141 | RTPRN-STG-9 | TCTACCGGTCACATGATCCTGGCTTAC |
| 142 | RTPRN-FGD-9 | TTCGGTGACCACCCGGGTCACTCTTAC |
| 143 | RTPRN-IST-10 | ATCTCTACCGGTCACATGATCCTGGCTTAC |
| 144 | RTPRN-FQD-8 | TTCCAGGACTCTGGTCTGCTGTAC |
| 145 | RTPRN-QLF-10 | CAGCTGTTCCAGGACTCTGGTCTGCTGTAC |
| 146 | RTPRN-LSW-10 | CTGTCTTGGCACGACGACCTGACCCAGTAC |
| 147 | RTPRN-WPD-9 | TGGCCGGACGAAGGTGCTTCTCTGTAC |
| 148 | GFAP-ALD-9 | GCTCTGGACATCGAAATCGCTACCTAC |
| 149 | GFAP-LAL-10 | CTGGCTCTGGACATCGAAATCGCTACCTAC |
| 150 | PPI-42-51 | GTTTGCGGTGAACGTGGTTTCTTCTACACC |
| 151 | PPI-44-51 | GGTGAACGTGGTTTCTTCTACACC |
| 152 | PPI-41-50 | CTGGTTTGCGGTGAACGTGGTTTCTTCTAC |
| 153 | INS-15-25 | GCTCTGTGGGGTCCGGACCCGGCTGCTGCTTTC |
| 154 | NP44-52 | TGCACCGAACTGAAACTGTCTGACTAC |
| 155 | NP140-148 | CACTCTAACCTGAACGACGCTACCTAC |
| 156 | NP273-281 | AAATCTTGCCTGCCGGCTTGCGTTTAC |
| 157 | PB1-590-599 | CTGGTTTCTGACGGTGGTCCGAACCTGTAC |
| 158 | PB1-591-599 | GTTTCTGACGGTGGTCCGAACCTGTAC |
| 159 | M1-ALA | GCTCTGGCTTCTTGCATGGGTCTGATCTAC |
| 160 | Gag71-79 | GGTTCTGAAGAACTGCGTTCTCTGTAC |
| 161 | Gag293-301 | TTCCGTGACTACGTTGACCGTTTCTAC |
| 162 | Pol63-71 | CAGCGTCCGCTGGTTACCATCAAAATC |
| 163 | Rev55-63 | ATCTCTGAACGTATCCTGTCTACCTAC |
| 164 | Gp160-787-795 | CGTCGTGGTTGGGAAGTTCTGAAATAC |
| 165 | INS-MAL-10 | ATGGCTCTGTGGATGCGTCTGCTGCCGCTG |
| 166 | INS-WMR-10 | TGGATGCGTCTGCTGCCGCTGCTGGCTCTG |
| 167 | INS-WMR-8 | TGGATGCGTCTGCTGCCGCTGCTG |
| 168 | INS-LWM-8 | CTGTGGATGCGTCTGCTGCCGCTG |
| 169 | INS-SLQ-8 | TCTCTGCAGAAACGTGGTATCGTT |
| 170 | INS-MAL-8 | ATGGCTCTGTGGATGCGTCTGCTG |
| 171 | GAD2-MMI-8 | ATGATGATCGCTCGTTTCAAAATG |
| 172 | GAD2-MMI-9 | ATGATGATCGCTCGTTTCAAAATGTTC |
| 173 | GAD2-MSR-9 | ATGTCTCGTAAACACAAATGGAAACTG |
| 174 | GAD2-LMS-10 | CTGATGTCTCGTAAACACAAATGGAAACTG |
| 175 | GAD2-SLK-9 | TCTCTGAAAAAAGGTGCTGCTGCTCTG |
| 176 | GAD2-FSL-10 | TTCTCTCTGAAAAAAGGTGCTGCTGCTCTG |
| 177 | GAD2-HPR-8 | CACCCGCGTTACTTCAACCAGCTG |
| 178 | GAD2-LMH-8 | CTGATGCACTGCCAGACCACCCTG |
| 179 | GAD2-AMM-10 | GCTATGATGATCGCTCGTTTCAAAATGTTC |
| 180 | GAD2-MSR-10 | ATGTCTCGTCTGTCTAAAGTTGCTCCGGTT |
| 181 | GAD2-MAA-10 | ATGGCTGCTCTGCCGCGTCTGATCGCTTTC |
| 182 | GAD2-MIA-8 | ATGATCGCTCGTTTCAAAATGTTC |
| 183 | GAD2-TLK-8 | ACCCTGAAAAAAATGCGTGAAATC |
| 184 | GAD2-EAK-10 | GAAGCTAAACAGAAAGGTTTCGTTCCGTTC |
| 185 | GAD2-RMM-9 | CGTATGATGGAATACGGTACCACCATG |

| | | |
|---|---|---|
| 186 | GAD2-EVK-10 | GAAGTTAAAGAAAAAGGTATGGCTGCTCTG |
| 187 | GAD2-EVK-9 | GAAGTTAAAGAAAAAGGTATGGCTGCT |
| 188 | GAD2-YAM-10 | TACGCTATGATGATCGCTCGTTTCAAAATG |
| 189 | GAD2-NPH-10 | AACCCGCACAAAATGATGGGTGTTCCGCTG |
| 190 | GAD2-SRK-8 | TCTCGTAAACACAAATGGAAACTG |
| 191 | GAD2-FQQ-9 | TTCCAGCAGGACAAACACTACGACCTG |
| 192 | GAD2-YAF-9 | TACGCTTTCCTGCACGCTACCGACCTG |
| 193 | GAD2-FSL-9 | TTCTCTCTGAAAAAAGGTGCTGCTGCT |
| 194 | GAD2-SLK-8 | TCTCTGAAAAAAGGTGCTGCTGCT |
| 195 | GAD2-TLK-9 | ACCCTGAAAAAAATGCGTGAAATCATC |
| 196 | GAD2-ERM-9 | GAACGTATGTCTCGTCTGTCTAAAGTT |
| 197 | ZNT8-YAK-9 | TACGCTAAATGGAAACTGTGCTCTGCT |
| 198 | ZNT8-AAK-9 | GCTGCTAAAATGTACGCTTTCACCCTG |
| 199 | ZNT8-CPR-9 | TGCCCGCGTGAACGTCCGGAAGAACTG |
| 200 | ZNT8-YAY-8 | TACGCTTACGCTAAATGGAAACTG |
| 201 | ZNT8-FLL-8 | TTCCTGCTGTCTCTGTTCTCTCTG |
| 202 | ZNT8-SVR-10 | TCTGTTCGTGCTGCTTTCGTTCACGCTCTG |
| 203 | ZNT8-NAS-8 | AACGCTTCTGTTCGTGCTGCTTTC |
| 204 | ZNT8-HSL-8 | CACTCTCTGCACATCTGGTCTCTG |
| 205 | IAPP-EVL-8 | GAAGTTCTGAAACGTGAACCGCTG |
| 206 | IAPP-LNH-9 | CTGAACCACCTGAAAGCTACCCCGATC |
| 207 | IAPP-ILK-8 | ATCCTGAAACTGCAGGTTTTCCTG |
| 208 | IAPP-MGI-9 | ATGGGTATCCTGAAACTGCAGGTTTTC |
| 209 | IAPP-MGI-10 | ATGGGTATCCTGAAACTGCAGGTTTTCCTG |
| 210 | IAPP-NTY-9 | AACACCTACGGTAAACGTAACGCTGTT |
| 211 | G6PC2-FLH-8 | TTCCTGCACCGTAACGGTGTTCTG |
| 212 | G6PC2-YLK-8 | TACCTGAAAACCAACCTGTTCCTG |
| 213 | G6PC2-NLI-8 | AACCTGATCTTCAAATGGATCCTG |
| 214 | G6PC2-YVM-8 | TACGTTATGGTTACCGCTGCTCTG |
| 215 | G6PC2-TLS-10 | ACCCTGTCTTTCCGTCTGCTGTGCGCTCTG |
| 216 | G6PC2-YLK-10 | TACCTGAAAACCAACCTGTTCCTGTTCCTG |
| 217 | G6PC2-YLK-9 | TACCTGAAAACCAACCTGTTCCTGTTC |
| 218 | G6PC2-SFR-8 | TCTTTCCGTCTGCTGTGCGCTCTG |
| 219 | G6PC2-TLH-9 | ACCCTGCACCGTCTGACCTGGTCTTTC |
| 220 | G6PC2-CGM-10 | TGCGGTATGGACAAATTCTCTATCACCCTG |
| 221 | G6PC2-CGM-8 | TGCGGTATGGACAAATTCTCTATC |
| 222 | G6PC2-NLI-9 | AACCTGATCTTCAAATGGAAATCTATC |
| 223 | G6PC2-WPC-10 | TGGCCGTGCAACGGTCGTATCCTGTGCCTG |
| 224 | PTPRN-VLL-9 | GTTCTGCTGGAAAAAAAAATCTCCGCTG |
| 225 | PTPRN-FLV-8 | TTCCTGGTTCGTTCTTTCTACCTG |
| 226 | PTPRN-HLR-8 | CACCTGCGTAACCGTGACCGTCTG |
| 227 | PTPRN-SPM-8 | TCTCCGATGCGTTCTGTTCTGCTG |
| 228 | PTPRN-AAL-9 | GCTGCTCTGCAGCGTCTGGCTGCTGTT |
| 229 | PTPRN-LPA-8 | CTGCCGGCTCGTACCTCTCCGATG |
| 230 | PTPRN-LLE-8 | CTGCTGGAAAAAAAAATCTCCGCTG |
| 231 | PTPRN-DKE-8 | GACAAAGAACGTCTGGCTGCTCTG |
| 232 | PTPRN-HAR-8 | CACGCTCGTATCAAACTGAAAGTT |

| | | |
|---|---|---|
| 233 | PTPRN-YRG-8 | TACCGTGGTCGTTCTTGCCCGATC |
| 234 | PTPRN-QQD-10 | CAGCAGGACAAAGAACGTCTGGCTGCTCTG |
| 235 | PTPRN-ELP-9 | GAACTGCCGGCTCGTACCTCTCCGATG |
| 236 | PTPRN-CYR-9 | TGCTACCGTGGTCGTTCTTGCCCGATC |
| 237 | PTPRN-RPR-8 | CGTCCGCGTGACCGTTCTGGTCTG |
| 238 | PTPRN-SPM-10 | TCTCCGATGCGTTCTGTTCTGCTGACCCTG |
| 239 | PTPRN-ALQ-8 | GCTCTGCAGCGTCTGGCTGCTGTT |
| 240 | PTPRN-AAL-10 | GCTGCTCTGCAGCGTCTGGCTGCTGTTCTG |
| 241 | PTPRN-HLR-9 | CACCTGCGTAACCGTGACCGTCTGGCT |
| 242 | PTPRN-LAK-8 | CTGGCTAAAGAATGGCAGGCTCTG |
| 243 | PTPRN-QDK-9 | CAGGACAAAGAACGTCTGGCTGCTCTG |
| 244 | GFAP-NLQ-9 | AACCTGCAGATCCGTGAAACCTCTCTG |
| 245 | GFAP-LLN-8 | CTGCTGAACGTTAAACTGGCTCTG |
| 246 | GFAP-ELRL-9 | GAACTGCGTCTGCGTCTGGACCAGCTG |
| 247 | GFAP-MER-9 | ATGGAACGTCGTCGTATCACCTCTGCT |
| 248 | GFAP-WYR-9 | TGGTACCGTTCTAAATTCGCTGACCTG |
| 249 | GFAP-HLK-8 | CACCTGAAACGTAACATCGTTGTT |
| 250 | GFAP-YRR-8 | TACCGTCGTCAGCTGCAGTCTCTG |
| 251 | GFAP-YRS-8 | TACCGTTCTAAATTCGCTGACCTG |
| 252 | GFAP-SNL-10 | TCTAACCTGCAGATCCGTGAAACCTCTCTG |
| 253 | GFAP-ELRE-9 | GAACTGCGTGAACTGCGTCTGCGTCTG |
| 254 | GFAP-DYR-9 | GACTACCGTCGTCAGCTGCAGTCTCTG |
| 255 | GFAP-SAA-8 | TCTGCTGCTCGTCGTTCTTACGTT |
| 256 | GFAP-EGH-9 | GAAGGTCACCTGAAACGTAACATCGTT |
| 257 | GFAP-MER-10 | ATGGAACGTCGTCGTATCACCTCTGCTGCT |
| 258 | GFAP-LRL-8 | CTGCGTCTGCGTCTGGACCAGCTG |
| 259 | GFAP-DLE-9 | GACCTGGAACGTAAAATCGAATCTCTG |
| 260 | GFAP-LQI-8 | CTGCAGATCCGTGAAACCTCTCTG |
| 261 | GFAP-REL-10 | CGTGAACTGCGTCTGCGTCTGGACCAGCTG |
| 262 | GFAP-LAR-8 | CTGGCTCGTATGCCGCCGCCGCTG |
| 263 | GFAP-EIR-9 | GAAATCCGTACCCAGTACGAAGCTATG |
| 264 | GFAP-GPG-8 | GGTCCGGGTACCCGTCTGTCTCTG |
| 265 | NP263-271 | GCTGACCGTGGTCTGCTGCGTGACATC |
| 266 | NP413-421 | GCTCTGAAATGCAAAGGTTTCCACGTT |
| 267 | NP380-388 | GAACTGCGTTCTCGTTACTGGGCTATC |
| 268 | NP225-233 | ATCCTGAAAGGTAAATTCCAGACCGCT |
| 269 | NP30-38 | CGTCCGATCATCCGTCCGGCTACCCTG |
| 270 | Gag74-82 | GAACTGCGTTCTCTGTACAACACCGTT |
| 271 | Gag260-267 | GAAATCTACAAACGTTGGATCATC |
| 272 | gp160-2-10 | CGTGTTAAAGAAAAATACCAGCACCTG |
| 273 | gp160-586-593 | TACCTGAAAGACCAGCAGCTGCTG |
| 274 | Nef13-20 | TGGCCGACCGTTCGTGAACGTATG |
| 275 | Nef90-97 | TTCCTGAAAGAAAAAGGTGGTCTG |
| 276 | Pol173-181 | GGTCCGAAAGTTAAACAGTGGCCGCTG |
| 277 | EBNA3A-193-201 | TTCCTGCGTGGTCGTGCTTACGGTCTG |
| 278 | BZLF1-190-197 | CGTGCTAAATTCAAACAGCTGCTG |
| 279 | EBNA3A-158-166 | CAGGCTAAATGGCGTCTGCAGACCCTG |

| | | |
|---|---|---|
| 280 | LMP2A-CPL | TGCCCGCTGTCTAAAATCCTGCTG |
| 281* | CMV-empty | AACCTGGTTCCGATGGTTGCTACCGTT |
| 282* | YFV-empty | CTGCTGTGGAACGGTCCGATGGCTGTT |

*Empty: Negative control without antigen loaded on pMHC.

nents with primary CD8+ T cells

| AA sequence of antigen | Gene | Fluorescence label | HLA | Group |
|---|---|---|---|---|
| RQFGPDWIVA | ADH | PE | A0201 | T1D |
| MVWGPDPLYV | DUF5119 | PE | A0201 | T1D |
| NLAQDLATV | GFAP | PE | A0201 | T1D |
| QLARQQVHV | GFAP | PE | A0201 | T1D |
| FLQDVMNIL | GAD65 | PE | A0201 | T1D |
| LLQEYNWEL | GAD65 | PE | A0201 | T1D |
| RMMEYGTTMV | GAD65 | PE | A0201 | T1D |
| VMNILLQYVV | GAD65 | PE | A0201 | T1D |
| VMNILLQYV | GAD65 | PE | A0201 | T1D |
| ELAEYLYNI | GAD65 | PE | A0201 | T1D |
| ILMHCQTTL | GAD65 | PE | A0201 | T1D |
| MLYQHLLPL | INS | PE | A0201 | T1D |
| GIVEQCCTSI | INS | PE | A0201 | T1D |
| ALWMRLLPLL | INS | PE | A0201 | T1D |
| LALWGPDPAA | INS | PE | A0201 | T1D |
| RLLPLLALLAL | INS | PE | A0201 | T1D |
| ALWMRLLPL | INS | PE | A0201 | T1D |
| HLVEALYLV | INS | PE | A0201 | T1D |
| SLQKRGIVEQ | INS | PE | A0201 | T1D |
| SLQPLALEG | INS | PE | A0201 | T1D |
| SLYQLENYC | INS | PE | A0201 | T1D |
| VCGERGFFYT | INS | PE | A0201 | T1D |
| ALWGPDPAAA | INS | PE | A0201 | T1D |
| RLLPLLALL | INS | PE | A0201 | T1D |
| WGPDPAAA | INS | PE | A0201 | T1D |
| FLIVLSVAL | IAPP | PE | A0201 | T1D |
| KLQVFLIVL | IAPP | PE | A0201 | T1D |
| FLWSVFMLI | IGRP | PE | A0201 | T1D |
| FLFAVGFYL | IGRP | PE | A0201 | T1D |
| LNIDLLWSV | IGRP | PE | A0201 | T1D |
| VLFGLGFAI | IGRP | PE | A0201 | T1D |
| FLWSVFWLI | IGRP | PE | A0201 | T1D |
| NLFLFLFAV | IGRP | PE | A0201 | T1D |
| YLLLRVLNI | IGRP | PE | A0201 | T1D |
| HLCGSHLVEA | INS | PE | A0201 | T1D |
| SHLVEALYLV | INS | PE | A0201 | T1D |
| LCGSHLVEAL | INS | PE | A0201 | T1D |
| ALTAVAEEV | PTPRN | PE | A0201 | T1D |
| SLYHVYEVNL | PTPRN | PE | A0201 | T1D |
| TIADFWQMV | PTPRN | PE | A0201 | T1D |
| VIVMLTPLV | PTPRN | PE | A0201 | T1D |
| LLPPLLEHL | PTPRN | PE | A0201 | T1D |
| SLAAGVKLL | PTPRN | PE | A0201 | T1D |
| SLSPLQAEL | PTPRN | PE | A0201 | T1D |

| | | | | |
|---|---|---|---|---|
| MVWESGCTV | PTPRN | PE | A0201 | T1D |
| VMIIVSSLAV | ZNT8 | PE | A0201 | T1D |
| ALGDLFQSI | ZNT8 | PE | A0201 | T1D |
| DLTSFLLSL | ZNT8 | PE | A0201 | T1D |
| EILGALLSI | ZNT8 | PE | A0201 | T1D |
| FLLSLFSLWL | ZNT8 | PE | A0201 | T1D |
| ILAVDGVLSV | ZNT8 | PE | A0201 | T1D |
| ILGALLSIL | ZNT8 | PE | A0201 | T1D |
| ILKDFSILL | ZNT8 | PE | A0201 | T1D |
| ILSAHVATA | ZNT8 | PE | A0201 | T1D |
| LLIDLTSFL | ZNT8 | PE | A0201 | T1D |
| LLMEGVPKSL | ZNT8 | PE | A0201 | T1D |
| SISVLISAL | ZNT8 | PE | A0201 | T1D |
| SLNYSGVKEL | ZNT8 | PE | A0201 | T1D |
| SVHSLHIWSL | ZNT8 | PE | A0201 | T1D |
| VVTGVLVYL | ZNT8 | PE | A0201 | T1D |
| FIFSILVLA | ZNT8 | PE | A0201 | T1D |
| IQATVMIIV | ZNT8 | PE | A0201 | T1D |
| KMYAFTLES | ZNT8 | PE | A0201 | T1D |
| KSLNYSGVK | ZNT8 | PE | A0201 | T1D |
| LAVDGVLSV | ZNT8 | PE | A0201 | T1D |
| LLSLFSLWL | ZNT8 | PE | A0201 | T1D |
| RLLYPDYQI | ZNT8 | PE | A0201 | T1D |
| TMHSLTIQM | ZNT8 | PE | A0201 | T1D |
| VAANIVLTV | ZNT8 | PE | A0201 | T1D |
| CLGHNHKEV | ZNT8 | PE | A0201 | T1D |
| KIADPICTFI | ZNT8 | PE | A0201 | T1D |
| KMYAFTLESV | ZNT8 | PE | A0201 | T1D |
| LLIDLTSFLL | ZNT8 | PE | A0201 | T1D |
| LLSILCIWV | ZNT8 | PE | A0201 | T1D |
| SLYNTVATLY | HIV | APC | A0201 | Viral |
| FLGKIWPSYK | HIV | APC | A0201 | Viral |
| LVGPTPVNI | HIV | APC | A0201 | Viral |
| ALVEICTEM | HIV | APC | A0201 | Viral |
| VIYQYMDDL | HIV | APC | A0201 | Viral |
| ILKEPVHGV | HIV | APC | A0201 | Viral |
| AIIRILQQL | HIV | APC | A0201 | Viral |
| RGPGRAFVTI | HIV | APC | A0201 | Viral |
| SLLNATDIAV | HIV | APC | A0201 | Viral |
| LLNATDIAV | HIV | APC | A0201 | Viral |
| PLTFGWCYKL | FLU | APC | A0201 | Viral |
| VLEWRFDSRL | FLU | APC | A0201 | Viral |
| GILGFVFTL | FLU | APC | A0201 | Viral |
| KLYQNPTTYI | FLU | APC | A0201 | Viral |
| RLYQNPTTYI | FLU | APC | A0201 | Viral |
| AIMDKNIIL | FLU | APC | A0201 | Viral |
| FMYSDFHFI | FLU | APC | A0201 | Viral |

| | | | | |
|---|---|---|---|---|
| KLVALGINAV | HCV | APC | A0201 | Viral |
| LLFNILGGWV | HCV | APC | A0201 | Viral |
| CINGVCWTV | HCV | APC | A0201 | Viral |
| YLLPRRGPRL | HCV | APC | A0201 | Viral |
| CLGGLLTMV | EBV | APC | A0201 | Viral |
| YLQQNWWTL | EBV | APC | A0201 | Viral |
| YLLEMLWRL | EBV | APC | A0201 | Viral |
| YVLDHLIVV | EBV | APC | A0201 | Viral |
| GLCTLVAML | EBV | APC | A0201 | Viral |
| FLYALALLL | EBV | APC | A0201 | Viral |
| YLLPGWKL | ROTA | APC | A0201 | Viral |
| SLISGMWLL | ROTA | APC | A0201 | Viral |
| TLLANVTAV | ROTA | APC | A0201 | Viral |
| EVKEKHEFL | CVB | APC | A0201 | Viral |
| ILMNDQEVGV | CVB | APC | A0201 | Viral |
| GIIYIIYKL | CVB | APC | A0201 | Viral |
| EAAGIGILTV | MART1 | PE | A0201 | self |
| ELAGIGILTV | MART1 | PE | A0201 | self |
| ALAGIGILTV | MART1 | PE | A0201 | self |
| AAGIGILTV | MART1 | PE | A0201 | self |
| ALGIGILTV | MART1 | PE | A0201 | self |
| LLAGIGTVPI | PGT | PE | A0201 | self |
| YLVALGINAV | HCV | APC | A0201 | Viral |
| YLVALGVNAV | HCV | APC | A0201 | Viral |
| KLVALGINNV | HCV | APC | A0201 | Viral |
| SLVALGINAV | HCV | APC | A0201 | Viral |
| KIVALGINAV | HCV | APC | A0201 | Viral |
| CTSICSLY | INS | PE | A0101 | T1D |
| CGSHLVEALY | INS | PE | A0101 | T1D |
| GSHLVEALY | INS | PE | A0101 | T1D |
| CLELAEYLY | GAD65 | PE | A0101 | T1D |
| STANTNMFTY | GAD65 | PE | A0101 | T1D |
| KCLELAEYLY | GAD65 | PE | A0101 | T1D |
| QQDKHYDLSY | GAD65 | PE | A0101 | T1D |
| VSATAGTTVY | GAD65 | PE | A0101 | T1D |
| STKVIDFHY | GAD65 | PE | A0101 | T1D |
| YLACERLLY | ZNT8 | PE | A0101 | T1D |
| VTDAAHLLI | ZNT8 | PE | A0101 | T1D |
| PTEKGANEY | ZNT8 | PE | A0101 | T1D |
| LIDLTSFLL | ZNT8 | PE | A0101 | T1D |
| KPTEKGANEY | ZNT8 | PE | A0101 | T1D |
| VVTDAAHLLI | ZNT8 | PE | A0101 | T1D |
| LTSFLLSLF | ZNT8 | PE | A0101 | T1D |
| STNVGSNTY | IAPP | PE | A0101 | T1D |
| SSTNVGSNTY | IAPP | PE | A0101 | T1D |
| LTSLTILQLY | IGRP | PE | A0101 | T1D |
| PTHEEHLFY | IGRP | PE | A0101 | T1D |

| | | | | |
|---|---|---|---|---|
| IPTHEEHLFY | IGRP | PE | A0101 | T1D |
| TSLTILQLY | IGRP | PE | A0101 | T1D |
| STGHMILAY | PTPRN | PE | A0101 | T1D |
| FGDHPGHSY | PTPRN | PE | A0101 | T1D |
| ISTGHMILAY | PTPRN | PE | A0101 | T1D |
| FQDSGLLY | PTPRN | PE | A0101 | T1D |
| QLFQDSGLLY | PTPRN | PE | A0101 | T1D |
| LSWHDDLTQY | PTPRN | PE | A0101 | T1D |
| WPDEGASLY | PTPRN | PE | A0101 | T1D |
| ALDIEIATY | GFAP | PE | A0101 | T1D |
| LALDIEIATY | GFAP | PE | A0101 | T1D |
| VCGERGFFYT | INS | PE | A0101 | T1D |
| GERGFFYT | INS | PE | A0101 | T1D |
| LVCGERGFFY | INS | PE | A0101 | T1D |
| ALWGPDPAAAF | INS | PE | A0101 | T1D |
| CTELKLSDY | FLU | APC | A0101 | Viral |
| HSNLNDATY | FLU | APC | A0101 | Viral |
| KSCLPACVY | FLU | APC | A0101 | Viral |
| LVSDGGPNLY | FLU | APC | A0101 | Viral |
| VSDGGPNLY | FLU | APC | A0101 | Viral |
| ALASCMGLIY | FLU | APC | A0101 | Viral |
| GSEELRSLY | HIV | APC | A0101 | Viral |
| FRDYVDRFY | HIV | APC | A0101 | Viral |
| QRPLVTIKI | HIV | APC | A0101 | Viral |
| ISERILSTY | HIV | APC | A0101 | Viral |
| RRGWEVLKY | HIV | APC | A0101 | Viral |
| MALWMRLLPL | INS | PE | B0801 | T1D |
| WMRLLPLLAL | INS | PE | B0801 | T1D |
| WMRLLPLL | INS | PE | B0801 | T1D |
| LWMRLLPL | INS | PE | B0801 | T1D |
| SLQKRGIV | INS | PE | B0801 | T1D |
| MALWMRLL | INS | PE | B0801 | T1D |
| MMIARFKM | GAD65 | PE | B0801 | T1D |
| MMIARFKMF | GAD65 | PE | B0801 | T1D |
| MSRKHKWKL | GAD65 | PE | B0801 | T1D |
| LMSRKHKWKL | GAD65 | PE | B0801 | T1D |
| SLKKGAAAL | GAD65 | PE | B0801 | T1D |
| FSLKKGAAAL | GAD65 | PE | B0801 | T1D |
| HPRYFNQL | GAD65 | PE | B0801 | T1D |
| LMHCQTTL | GAD65 | PE | B0801 | T1D |
| AMMIARFKMF | GAD65 | PE | B0801 | T1D |
| MSRLSKVAPV | GAD65 | PE | B0801 | T1D |
| MAALPRLIAF | GAD65 | PE | B0801 | T1D |
| MIARFKMF | GAD65 | PE | B0801 | T1D |
| TLKKMREI | GAD65 | PE | B0801 | T1D |
| EAKQKGFVPF | GAD65 | PE | B0801 | T1D |
| RMMEYGTTM | GAD65 | PE | B0801 | T1D |

| | | | | |
|---|---|---|---|---|
| EVKEKGMAAL | GAD65 | PE | B0801 | T1D |
| EVKEKGMAA | GAD65 | PE | B0801 | T1D |
| YAMMIARFKM | GAD65 | PE | B0801 | T1D |
| NPHKMMGVPL | GAD65 | PE | B0801 | T1D |
| SRKHKWKL | GAD65 | PE | B0801 | T1D |
| FQQDKHYDL | GAD65 | PE | B0801 | T1D |
| YAFLHATDL | GAD65 | PE | B0801 | T1D |
| FSLKKGAAA | GAD65 | PE | B0801 | T1D |
| SLKKGAAA | GAD65 | PE | B0801 | T1D |
| TLKKMREII | GAD65 | PE | B0801 | T1D |
| ERMSRLSKV | GAD65 | PE | B0801 | T1D |
| YAKWKLCSA | ZNT8 | PE | B0801 | T1D |
| AAKMYAFTL | ZNT8 | PE | B0801 | T1D |
| CPRERPEEL | ZNT8 | PE | B0801 | T1D |
| YAYAKWKL | ZNT8 | PE | B0801 | T1D |
| FLLSLFSL | ZNT8 | PE | B0801 | T1D |
| SVRAAFVHAL | ZNT8 | PE | B0801 | T1D |
| NASVRAAF | ZNT8 | PE | B0801 | T1D |
| HSLHIWSL | ZNT8 | PE | B0801 | T1D |
| EVLKREPL | IAPP | PE | B0801 | T1D |
| LNHLKATPI | IAPP | PE | B0801 | T1D |
| ILKLQVFL | IAPP | PE | B0801 | T1D |
| MGILKLQVF | IAPP | PE | B0801 | T1D |
| MGILKLQVFL | IAPP | PE | B0801 | T1D |
| NTYGKRNAV | IAPP | PE | B0801 | T1D |
| FLHRNGVL | IGRP | PE | B0801 | T1D |
| YLKTNLFL | IGRP | PE | B0801 | T1D |
| NLIFKWIL | IGRP | PE | B0801 | T1D |
| YVMVTAAL | IGRP | PE | B0801 | T1D |
| TLSFRLLCAL | IGRP | PE | B0801 | T1D |
| YLKTNLFLFL | IGRP | PE | B0801 | T1D |
| YLKTNLFLF | IGRP | PE | B0801 | T1D |
| SFRLLCAL | IGRP | PE | B0801 | T1D |
| TLHRLTWSF | IGRP | PE | B0801 | T1D |
| CGMDKFSITL | IGRP | PE | B0801 | T1D |
| CGMDKFSI | IGRP | PE | B0801 | T1D |
| NLIFKWKSI | IGRP | PE | B0801 | T1D |
| WPCNGRILCL | IGRP | PE | B0801 | T1D |
| VLLEKKSPL | PTPRN | PE | B0801 | T1D |
| FLVRSFYL | PTPRN | PE | B0801 | T1D |
| HLRNRDRL | PTPRN | PE | B0801 | T1D |
| SPMRSVLL | PTPRN | PE | B0801 | T1D |
| AALQRLAAV | PTPRN | PE | B0801 | T1D |
| LPARTSPM | PTPRN | PE | B0801 | T1D |
| LLEKKSPL | PTPRN | PE | B0801 | T1D |
| DKERLAAL | PTPRN | PE | B0801 | T1D |
| HARIKLKV | PTPRN | PE | B0801 | T1D |

| | | | | |
|---|---|---|---|---|
| YRGRSCPI | PTPRN | PE | B0801 | T1D |
| QQDKERLAAL | PTPRN | PE | B0801 | T1D |
| ELPARTSPM | PTPRN | PE | B0801 | T1D |
| CYRGRSCPI | PTPRN | PE | B0801 | T1D |
| RPRDRSGL | PTPRN | PE | B0801 | T1D |
| SPMRSVLLTL | PTPRN | PE | B0801 | T1D |
| ALQRLAAV | PTPRN | PE | B0801 | T1D |
| AALQRLAAVL | PTPRN | PE | B0801 | T1D |
| HLRNRDRLA | PTPRN | PE | B0801 | T1D |
| LAKEWQAL | PTPRN | PE | B0801 | T1D |
| QDKERLAAL | PTPRN | PE | B0801 | T1D |
| NLQIRETSL | GFAP | PE | B0801 | T1D |
| LLNVKLAL | GFAP | PE | B0801 | T1D |
| ELRLRLDQL | GFAP | PE | B0801 | T1D |
| MERRRITSA | GFAP | PE | B0801 | T1D |
| WYRSKFADL | GFAP | PE | B0801 | T1D |
| HLKRNIVV | GFAP | PE | B0801 | T1D |
| YRRQLQSL | GFAP | PE | B0801 | T1D |
| YRSKFADL | GFAP | PE | B0801 | T1D |
| SNLQIRETSL | GFAP | PE | B0801 | T1D |
| ELRELRLRL | GFAP | PE | B0801 | T1D |
| DYRRQLQSL | GFAP | PE | B0801 | T1D |
| SAARRSYV | GFAP | PE | B0801 | T1D |
| EGHLKRNIV | GFAP | PE | B0801 | T1D |
| MERRRITSAA | GFAP | PE | B0801 | T1D |
| LRLRLDQL | GFAP | PE | B0801 | T1D |
| DLERKIESL | GFAP | PE | B0801 | T1D |
| LQIRETSL | GFAP | PE | B0801 | T1D |
| RELRLRLDQL | GFAP | PE | B0801 | T1D |
| LARMPPPL | GFAP | PE | B0801 | T1D |
| EIRTQYEAM | GFAP | PE | B0801 | T1D |
| GPGTRLSL | GFAP | PE | B0801 | T1D |
| ADRGLLRDI | FLU | APC | B0801 | Viral |
| ALKCKGFHV | FLU | APC | B0801 | Viral |
| ELRSRYWAI | FLU | APC | B0801 | Viral |
| ILKGKFQTA | FLU | APC | B0801 | Viral |
| RPIIRPATL | FLU | APC | B0801 | Viral |
| ELRSLYNTV | HIV | APC | B0801 | Viral |
| EIYKRWII | HIV | APC | B0801 | Viral |
| RVKEKYQHL | HIV | APC | B0801 | Viral |
| YLKDQQLL | HIV | APC | B0801 | Viral |
| WPTVRERM | HIV | APC | B0801 | Viral |
| FLKEKGGL | HIV | APC | B0801 | Viral |
| GPKVKQWPL | HIV | APC | B0801 | Viral |
| FLRGRAYGL | EBV | APC | B0801 | Viral |
| RAKFKQLL | EBV | APC | B0801 | Viral |
| QAKWRLQTL | EBV | APC | B0801 | Viral |

| | | | | |
|---|---|---|---|---|
| CPLSKILL | EBV | APC | B0801 | Viral |
| Empty | Empty | PE | A0201 | Empty |
| Empty | Empty | APC | A0201 | Empty |

**Supplementary Table 5: non-T1D healthy subjects, T1D and T2D subjects used in the TetTCR-Se**

| Donor | Chip | Disease Status | Sex | Age | Auto-antibody test |
|---|---|---|---|---|---|
| Clones | Chip1 | N/A | N/A | N/A | N/A |
| T1D_1 | Chip2 | T1D | Male | 23 | GAD-, IA2-, ZnT8- |
| T1D_2 | Chip2 | T1D | Male | 40 | GAD+, IA2-, ZnT8+ |
| T1D_3 | Chip2 | T1D | Male | 27 | GAD-, IA2-, ZnT8- |
| healthy_1 | Chip2 | healthy | Male | 26 | N/A |
| healthy_2 | Chip2 | healthy | Female | 34 | N/A |
| T1D_4 | Chip3 | T1D | Female | 27 | GAD-, IA2-, ZnT8+ |
| T1D_5 | Chip3 | T1D | Female | 21 | GAD+, IA2-, ZnT8- |
| T1D_6 | Chip3 | T1D | Male | 48 | GAD+, IA2-, ZnT8- |
| healthy_3 | Chip3 | healthy | Female | 46 | N/A |
| healthy_4 | Chip3 | healthy | Male | 59 | N/A |
| healthy_5 | Chip3 | healthy | Male | 27 | N/A |
| T1D_7 | Chip4 | T1D | Male | 38 | GAD+, IA2-, ZnT8- |
| T1D_8 | Chip4 | T1D | Male | 21 | GAD-, IA2+, ZnT8+ |
| healthy_6 | Chip5 | healthy | Male | 37 | N/A |
| healthy_7 | Chip5 | healthy | Female | 27 | N/A |
| healthy_8 | Chip5 | healthy | Female | 25 | N/A |
| healthy_9 | Chip5 | healthy | Female | 19 | N/A |
| healthy_10 | Chip5 | healthy | Female | 44 | N/A |
| T2D_1 | Chip6 | T2D, later diagnosed with Latent Autoimmune Diabetes in Adults | Male | 50 | GAD+ |

**eqHD experiments.**

| HLAs | SampleTag ID | TotalCD8 | Number of sorted foreign antigen specific cells | Number of sorted self antigen specific cells | Number of total sorted cells |
|---|---|---|---|---|---|
| A0201 | N/A | N/A | N/A | N/A | 10000 |
| A0201 | SampleTag6 | 751101 | 10000 | 4603 | 14603 |
| A0201 | SampleTag7 | 714919 | 7000 | 2289 | 9289 |
| A0101, B0801 | SampleTag8 | 271299 | 441 | 81 | 522 |
| A0201 | SampleTag9 | 166522 | 2400 | 961 | 3361 |
| A0101, A0201, B0801 | SampleTag10 | 103735 | 584 | 1093 | 1677 |
| A0101, A0201, B0801 | SampleTag6 | 326035 | 6037 | 2256 | 8293 |
| A0101, B0801 | SampleTag7 | 698189 | 8120 | 1595 | 9715 |
| A0101, A0201, B0801 | SampleTag8 | 598137 | 5300 | 3315 | 8615 |
| A0101, A0201 | SampleTag9 | 482681 | 3347 | 2564 | 5911 |
| A0201, B0801 | SampleTag10 | 250323 | 3246 | 914 | 4160 |
| A0101, B0801 | SampleTag11 | 196883 | 434 | 239 | 673 |
| A0201 | SampleTag2 | 434254 | 2674 | 2545 | 5219 |
| A0201 | SampleTag3 | 112918 | 815 | 732 | 1547 |
| A0201 | SampleTag11 | 192387 | 3242 | 787 | 4029 |
| A0201 | SampleTag10 | 196243 | 1117 | 1178 | 2295 |
| A0101, B0801 | SampleTag7 | 258144 | 2129 | 812 | 2941 |
| A0101, B0801 | SampleTag8 | 368229 | 777 | 621 | 1398 |
| A0101, B0801 | SampleTag9 | 45245 | 582 | 177 | 759 |
| A0201 | SampleTag5 | 178523 | 1105 | 1461 | 2566 |

| Number of cells recovered in Rhapsody | TCRa | TCRb | paired |
|---|---|---|---|
| 4533 | 3969(87.56%) | 4213(92.94%) | 3816(84.18%) |
| 5380 | 1406(26.13%) | 3181(59.13%) | 926(17.21%) |
| 3460 | 1294(37.40%) | 2594(74.97%) | 980(28.32%) |
| 171 | 32(18.71%) | 58(33.92%) | 9(5.26%) |
| 987 | 321(32.52%) | 538(54.51%) | 231(23.40%) |
| 463 | 113(24.41%) | 207(44.71%) | 73(15.77%) |
| 2897 | 1753(60.51%) | 2331(80.46%) | 1462(50.47%) |
| 3167 | 1856(58.60%) | 2565(80.99%) | 1570(49.57%) |
| 3239 | 1694(52.30%) | 2639(81.48%) | 1399(43.19%) |
| 1904 | 954(50.11%) | 1228(64.50%) | 656(34.45%) |
| 1210 | 400(33.06%) | 749(61.90%) | 273(22.56%) |
| 162 | 47(29.01%) | 83(51.23%) | 31(19.14%) |
| 1817 | 1083(59.60%) | 1356(74.63%) | 831(45.73%) |
| 549 | 381(69.40%) | 422(76.87%) | 302(55.01%) |
| 1247 | 470(37.69%) | 649(52.04%) | 266(21.33%) |
| 819 | 361(44.08%) | 554(67.64%) | 268(32.72%) |
| 1147 | 693(60.42%) | 696(60.68%) | 448(39.06%) |
| 436 | 133(30.50%) | 212(48.62%) | 69(15.83%) |
| 293 | 109(37.20%) | 187(63.82%) | 74(25.26%) |
| 725 | 236(32.55%) | 285(39.31%) | 147(20.28%) |

**Supplementary Table 6: Number of antigens and cells detected for non-T1D endogenous, T1D, and viral antigen**

| donor | group | Number of different antigens | Number of cells |
|---|---|---|---|
| healthy_1 | self | 6 | 163 |
| healthy_1 | T1D | 18 | 32 |
| healthy_1 | Viral | 22 | 510 |
| healthy_2 | self | 6 | 47 |
| healthy_2 | T1D | 19 | 29 |
| healthy_2 | Viral | 26 | 101 |
| healthy_3 | self | 6 | 290 |
| healthy_3 | T1D | 43 | 228 |
| healthy_3 | Viral | 29 | 676 |
| healthy_4 | self | 6 | 106 |
| healthy_4 | T1D | 31 | 41 |
| healthy_4 | Viral | 25 | 453 |
| healthy_5 | self | 0 | 0 |
| healthy_5 | T1D | 11 | 18 |
| healthy_5 | Viral | 11 | 56 |
| healthy_6 | self | 6 | 144 |
| healthy_6 | T1D | 26 | 50 |
| healthy_6 | Viral | 24 | 803 |
| healthy_7 | self | 6 | 252 |
| healthy_7 | T1D | 34 | 60 |
| healthy_7 | Viral | 28 | 173 |
| healthy_8 | self | 0 | 0 |
| healthy_8 | T1D | 52 | 234 |
| healthy_8 | Viral | 18 | 764 |
| healthy_9 | self | 0 | 0 |
| healthy_9 | T1D | 40 | 104 |
| healthy_9 | Viral | 19 | 197 |
| healthy_10 | self | 0 | 0 |
| healthy_10 | T1D | 18 | 22 |
| healthy_10 | Viral | 14 | 41 |
| T1D_1 | self | 6 | 830 |
| T1D_1 | T1D | 50 | 134 |
| T1D_1 | Viral | 31 | 2587 |
| T1D_2 | self | 6 | 345 |
| T1D_2 | T1D | 32 | 56 |
| T1D_2 | Viral | 21 | 1809 |
| T1D_3 | self | 0 | 0 |
| T1D_3 | T1D | 14 | 17 |
| T1D_3 | Viral | 7 | 105 |
| T1D_4 | self | 6 | 310 |
| T1D_4 | T1D | 65 | 148 |
| T1D_4 | Viral | 37 | 1349 |
| T1D_5 | self | 0 | 0 |

| | | | |
|---|---|---|---|
| T1D_5 | T1D | 56 | 278 |
| T1D_5 | Viral | 16 | 1206 |
| T1D_6 | self | 6 | 450 |
| T1D_6 | T1D | 80 | 305 |
| T1D_6 | Viral | 40 | 895 |
| T1D_7 | self | 6 | 549 |
| T1D_7 | T1D | 51 | 153 |
| T1D_7 | Viral | 29 | 605 |
| T1D_8 | self | 6 | 140 |
| T1D_8 | T1D | 27 | 43 |
| T1D_8 | Viral | 22 | 188 |

**Supplementary Table 7: Summary of transduced TCR sequences and their cognate antigen specifi**

| TCR_ID | Specificity | HLA | TCRaCDR3 |
|---|---|---|---|
| TCR051 | DUF5119-124-133\|INSDRIP-1-9\|PTPRN-797-805 | A0201 | CAYRSVPKVSGSRLTF |
| TCR054 | INS-WMR-8 | B0801 | CAVSGNTGKLIF |
| TCR055 | RTPRN-FGD-9 | A0101 | CAASLKTSYDKVIF |
| TCR060 | EBV-BLMF1 | A0201 | CAESTSGGKLIF |
| TCR062 | EBV-BLMF1 | A0201 | CVVNGKDSSYKLIF |

**cities**

| **TCRaTRAV** | **TCRbCDR3** | **TCRbTRAV** |
|---|---|---|
| TRAV38-2/DV8*01 | CASSLRGGQSYEQYF | TRBV12-4*01 |
| TRAV20*01 | CASSQWDNQPQHF | TRBV4-1*01 |
| TRAV23/DV6*01 | CAISEGAPNTEAFF | TRBV10-3*01 |
| TRAV5*01 | CSVGTGGTNEKLFF | TRBV29-1*01 |
| TRAV12-1*01 | CASSGGAVAPSEQFF | TRBV2*01 |

| Supplementary Table 8: |
|---|
| Name |
| DNA-linker |
| FLAG |
| FLAG_rxn2 |
| Peptide_barcodes* |
| IVTT_f |
| IVTT_r |

* peptide NT sequences a

**Oligonucleotide sequences used in TetTCR-SeqHD.**

| Sequence |
|---|
| C*T*TGTCGTCGTCGTCCATCAATGTAGATCAACCTCTGAGTACTA/iSP18//3AmC7/ |
| ATGGACGACGACGACAAG |
| GACGTGTGCTCTTCCGATCTATGGACGACGACGACAAG |
| ATGGACGACGACGACAAG [ peptide NT sequences ] TAACGAAGCACCTCGCTAAAAAAAAAAAAAAAAAAAAAAAAA |
| GCGAATTAATACGACTCACTATAGGGCTTAAGTATAAGGAGGAAAACATATGGACGACGACGACAAG |
| AAACCCCTCCGTTTAGAGAGGGGTTATGCTAGCGAGGTGCTTCGTTA |

are listed in **Supplementary Table 4**.
